# Multidimensional atlas of RNA-regulated proteins revealed by RNA-dependent thermal proteome profiling

**DOI:** 10.64898/2026.07.15.738841

**Authors:** Yongzuo Chen, Zeyu Liu, Yankun Wang, Wenjie Lu, Haoyuan Li, Wenjing Wang, Zhili Qiu, Yunjie Qiu, Hong Qing, Yixuan Xie, Nian Liu, Conggang Zhang, Ying Chen, Wei Qin

**Author notes:** These authors contributed equally (Yongzuo Chen, Zeyu Liu, Yankun Wang).

## Abstract

RNA and proteins interact pervasively in cellular processes, yet the functional relevance of most RNA-binding proteins remains unknown. To address this, we developed RNA-dependent thermal proteome profiling (RTPP), a method that identifies RNA-regulated proteins (RRPs) by detecting changes in a protein’s structural stability upon RNA binding, without requiring crosslinking or enrichment. Applying RTPP in HEK293T cells revealed 1,664 RRPs, including 257 previously undetected RNA-binding “orphans”. One such orphan, SGK3, binds the lncRNA CASC15, facilitating its PtdIns(3)P binding, endosomal recruitment and kinase activity. We further generated a tissue-specific RRP atlas, identifying interactions unique to particular organs, including hippocampus-specific RRPs dysregulated in Alzheimer’s disease. Integrating RTPP with proximity labeling resolved RNA-binding heterogeneities across subcellular compartments. Live-cell RNase treatment also uncovered proteins associated with cell surface RNA, revealing that RO60 form nanoscale clusters with glycoRNA. RTPP provides a powerful approach to decipher functional RNA-protein interactions across multiple biological dimensions.

## INTRODUCTION

Noncovalent RNA–protein interactions are prevalent in both transient and stable macromolecular complexes that underlie processes such as transcription, translation, and the stress response^1,2^. RNA-binding proteins (RBPs) play a pivotal role in regulating nearly every aspect of RNA function, including RNA processing, trafficking, and modifications^3–5^. Conversely, RNA also regulates the function of associated proteins; for instance, long non-coding RNAs can bind to and regulate a diverse range of proteins^6^. Defects in RNA-protein interactions are linked to a wide spectrum of severe diseases, ranging from cancer to neurodegenerative disorders^7^. Understanding how these two types of biomacromolecules impact on each other is critical to deciphering their physiological and pathological roles.

To gain a comprehensive understanding of the landscape of RBPs, numerous innovative strategies for large-scale RBP discovery have been reported in recent years^8^. One of these methods involves oligo(dT) bead capture, which enriches RBPs that bind to polyadenylated (poly(A)+) RNAs, leading to the identification of hundreds of poly(A)+ RBPs^9,10^. However, it’s worth noting that many RNA types and RNA in prokaryotic cells lack poly(A)+ tails. To address this, CARIC and RICK were developed based on metabolic labeling of RNA with alkynyl analogs, enabling the identification of poly(A)-independent RBPs^11,12^. Additionally, some research groups have reported organic-aqueous phase separation techniques for RBP enrichment, which are independent of RNA class and offer superior sensitivity^13–17^. Given the non-covalent and low-affinity nature of RNA-protein interactions, UV light was applied in all the aforementioned methods to crosslink RNA-protein complexes, preventing the loss of RBPs during the enrichment processes. While these methods have uncovered thousands of newly annotated RBPs, accounting for nearly one-third of the proteome, the majority of these proteins have unknown RNA-binding functions^18^. Some may randomly encounter RNA and get crosslinked, possessing negligible functional relationships with RNA. Moreover, UV crosslinking suffers from low efficiency and poor tissue penetration, limiting the application of these UV-based techniques for mapping RBPs *in vivo*^18–20^.

To identify RBPs that are functionally influenced by RNA, Caudron-Herger et al. developed R-DeeP, a crosslinking-free method for identifying RNA-dependent proteins^21^. R-DeeP employs sucrose-based density gradient ultracentrifugation to quantitatively analyze the impact of RNA depletion on protein interactomes. This approach complements previous RBP profiling methods and introduces the concept of “RNA dependence” to assign proteins whose complex formation is highly related to RNA, rather than being directly associated with RNA. This method has been further applied in mapping RNA-dependent proteins in the human malaria parasite, Plasmodium falciparum^22^. However, R-DeeP involves extensive fractionation of the proteome and complicated data analysis steps to determine altered protein complexes, limiting its applicability for discovering RNA-dependent proteins in tissues. Moreover, the impact of RNA depletion on protein function may not be directly revealed by protein interactomes, especially for proteins that do not interact with other proteins, or interact with other proteins in a transient and dynamic manner^23,24^.

In contrast to changes in protein interactomes, we propose that changes in protein structure should more directly reflect the functional status of a protein when its associated RNA is depleted. Thermal proteome profiling is a proteomic method developed for unbiased analysis of protein structural changes in living cells^25–32^, and more recently, in tissues^26^. Although it was initially designed for mapping drug targets, it has been extended to dissect the functional relevance of protein-protein interactions^31^ and post-translational modifications^27,32^. We introduce RTPP (RNA-dependent Thermal Proteome Profiling), a crosslinking- and enrichment-free proteomic method for identifying proteins that are *functionally* relevant to RNA, specifically RNA-regulated proteins (RRPs). We have established a streamlined workflow based on data-independent acquisition (DIA)^33^ for multiplexed quantification of melting curves, resulting in the identification of 1664 RRPs in human cells. One representative RNA-binding orphan, SGK3, binds the lncRNA CASC15 to facilitate its PtdIns(3)P binding, endosomal recruitment and kinase activity. Using this platform, we have created a comprehensive atlas of RRPs across primary organs in mice and explored RRPs related to Alzheimer’s disease (AD) in the mouse hippocampus. Additionally, we have integrated proximity labeling (PL) into this method to develop PL-RTPP for spatiotemporally resolved profiling of RRPs. We also developed cell surface RTPP to identify proteins associated with cell surface RNA, revealing nanoscale clusters of RRPs with glycoRNA.

## RESULTS

### Establishment of RTPP

Many RBPs interact with their RNA targets through molecular interactions involving chemical moieties between protein residues and RNA nucleotides^2^. Similar to how proteins interact with other molecules, such as drugs, RNA binding can induce structural changes in proteins and alter their thermal stability^25–32^. Indeed, a recent study found RNA ligands enhance the structural stability of their cognate binding proteins^34^. We posit that RBPs actively engaged with RNA are expected to display reduced thermal stability upon RNA digestion, whereas those with casual RNA interactions would exhibit minimal impact (**Figure 1A**). Notably, proteins that do not directly bind to RNAs but are closely associated with direct RBPs might also display altered thermal stability. For instance, a subset of components in RNA-binding complexes, such as ribosomes, do not make direct contact with RNA but rely on it for their functionality. Therefore, these proteins are also classified as RRPs.

**Figure 1.**
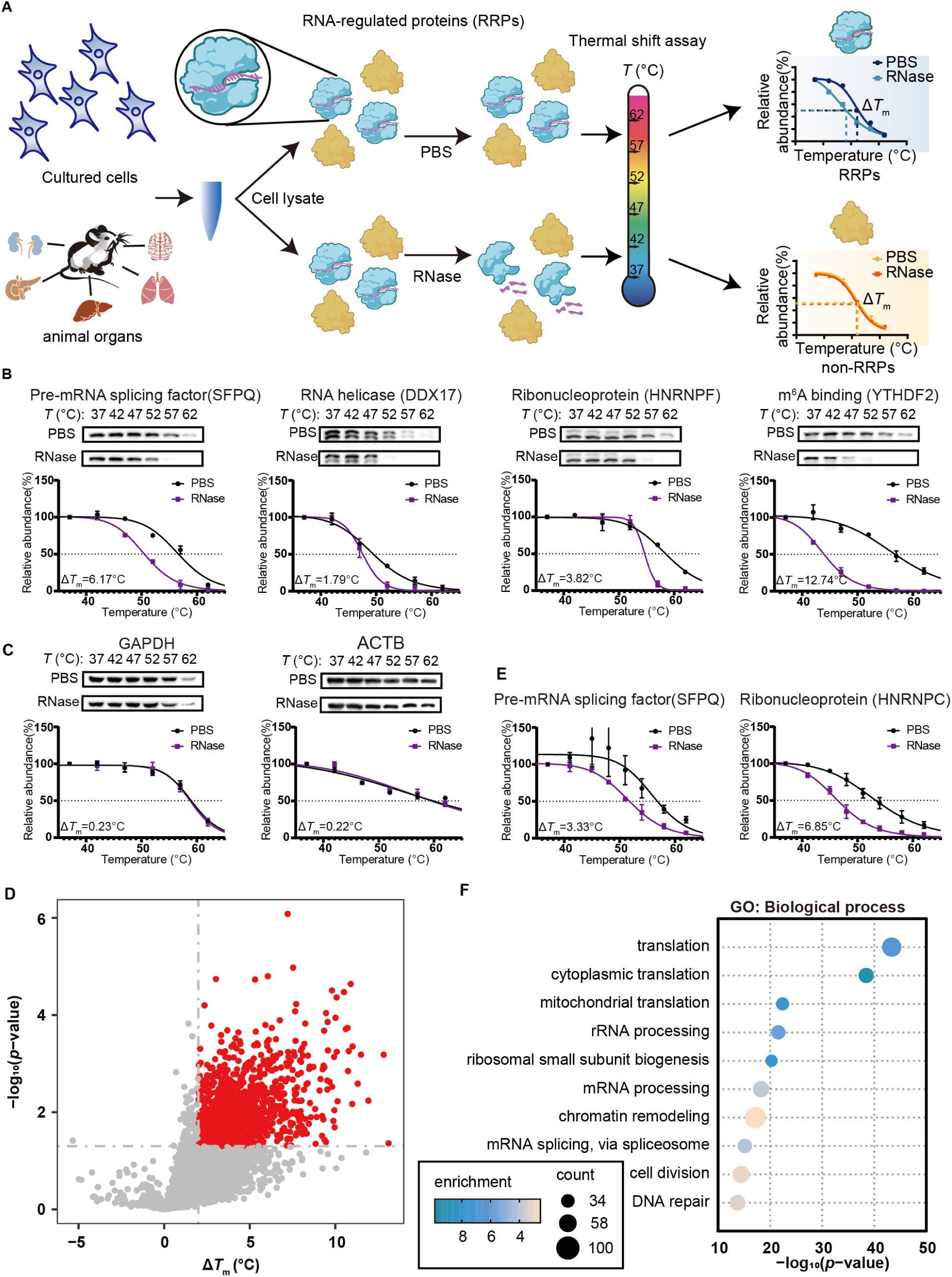
Development of RTPP for proteomic profiling of RNA-regulated proteins. (A) Schematic of the RTPP workflow. Lysates from cultured cells or tissue organs were treated separately with PBS and RNase, then divided into aliquots and heated at the indicated temperatures. The supernatants were analyzed by Western blotting or proteomic analysis. Protein intensity at each temperature point was measured to generate melting curves and calculate Δ*T*_m_. (B) Western blotting and thermal melting curves of four known RBPs: SFPQ, DDX17, HNRNPF and YTHDF2. (C) Western blotting and thermal melting curves of two known non-RBPs: GAPDH and ACTB. (D) Volcano plot of proteins quantified with Ä*T*_m_ in HEK293T cells using RTPP. Proteins with Δ*T*_m_ > 2°C and *p*-value < 0.05 were annotated as RRPs and labeled in red. The *p*-value was calculated with a two-sided student’s t-test. (E) Thermal melting curves of two known RBPs: SFPQ, HNRNPC. Error bars represent the mean ± s.d. (B, C, E) (F) GO-Biological Process analysis of RRPs identified in HEK293T cells.

To develop RTPP, we treat cell lysates with the same RNase cocktail used in R-DeeP^21^ to deplete RNA. The RNase-treated cell lysates are then incubated at various temperatures, ranging from 37°C to 62°C, for three minutes. After this incubation, we quantify the proteins in the remaining soluble fractions to determine melting curves. We compare the melting curves of the RNase-treated cell lysates against those of the control (cell lysates without RNase treatment) to quantify changes in protein melting temperature (Δ*T*_m_). The entire RTPP procedure is remarkably straightforward, requiring only a small amount of starting material (< 10^6^ cells) and a short processing time (< 2.5 hours) (**Figure S1A**). In contrast, enrichment-based RBP profiling methods, as well as R-DeeP, typically require much larger starting materials and longer processing times due to their labor-intensive sample preparation procedures^9–15,21^.

To evaluate the specificity of RTPP, we performed Western blotting on soluble fractions from HEK293T lysates, targeting a set of classical RBPs known to interact with distinct types of RNA. This set included the pre-mRNA splicing factor SFPQ (Splicing factor, proline- and glutamine-rich), RNA helicase DDX17 (Probable ATP-dependent RNA helicase), ribonucleoprotein HNRNPF (Heterogeneous nuclear ribonucleoprotein F), and m6A binding protein YTHDF2 (YTH domain-containing family protein 2). We observed that all four RBPs exhibited significantly decreased thermal stability, with Δ*T*_m_ values ranging from 1.79 to 12.74°C (**Figure 1B**). In contrast, we also probed the same samples for abundant non-RBPs such as ACTB and GAPDH, whose melting curves remained unchanged upon RNase treatment (**Figure 1C**). These results collectively indicate that RTPP can effectively discriminate RBPs from non-RBPs and offers a straightforward approach for verifying if a protein is RNA-related using Western blotting.

### Proteomic profiling of RNA-regulated proteins by RTPP

Following optimization and validation of RTPP, we performed proteomic analysis in HEK293T cells. Traditional thermal proteome profiling (TPP) relies on tandem mass tag (TMT) labeling and data-dependent acquisition (DDA), which limits throughput and suffers from stochastic data loss, complicating cross-experiment comparisons^25,28,29^. In contrast, we employed data-independent acquisition (DIA) for our RTPP analysis. DIA simultaneously fragments all peptides within predefined m/z windows, ensuring comprehensive coverage and superior quantification reproducibility. This powerful DIA-based pipeline enabled the robust and large-scale identification of RNA-regulated proteins (**Figure S1B**). The soluble fractions at each temperature point underwent the filter-aided sample preparation (FASP) procedure^35^, followed by Liquid chromatography–tandem mass spectrometry (LC-MS/MS) analysis in DIA mode.

The DIA data was analyzed using DIA-NN, a “library-free” approach that does not require spectral libraries generated by DDA methods^36^. Intensities at each temperature were ratioed to the intensity at 37°C, and melting curves were generated after data filtering and intensity normalization to calculate melting temperature (**Figure S1B**). In stark contrast to DDA, where replicate overlaps are relatively small, the vast majority of proteins were quantified in all three biological replicates of our DIA experiments (**Figure S1C**). Additionally, there was significant overlap in proteins quantified in both control and RNase-treated samples, with 4504 proteins exhibiting *T*_m_ values in both conditions (**Figure S1D**). The *T*_m_ values across replicates showed a high degree of correlation (**Figure S1E**), and the mean *T*_m_ values obtained from the control samples also correlated well with those from previously published datasets^27^ (**Figure S1F**), underscoring the high accuracy of DIA quantification for TPP experiments.

To define RRPs, we calculated Δ*T*_m_ (*T*_m_ ^control^ – *T*_m_ ^RNase^) for each quantified protein. Only proteins with average melting temperatures (*T*_m_) falling within the experimental temperature range were considered. We applied filtering criteria of Δ*T*_m_ > 2°C and a *p*-value < 0.05, consistent with established TPP protocols^25,29^ (**Table S1**). This approach identified 1143 RRPs exhibiting significantly reduced thermal stability upon RNA depletion (**Figure 1D**). In contrast, very few proteins showed increased stability following RNase treatment. This aligns with the premise that RNA binding enhances protein stability. As expected, established RBPs exhibited marked destabilization upon RNA loss (**Figure 1E**), whereas abundant non-RBPs displayed unchanged melting curves (**Figure S1G**), a result corroborated by Western blot validation. Several highly stable RBPs, such as SNRPD3 and PHF5A, were not captured by Δ*T*_m_ due to Tm values exceeding the experimental range, but showed clear RNase-dependent destabilization at multiple temperatures (**Figure S1H**). To include these, we performed single-temperature comparisons, adding 521 RRPs with significant destabilization at two or more points. In total, RTPP identified RRPs strongly enriched for RNA-related Gene Ontology (GO) terms, particularly in translation and rRNA processing (**Figure 1F, S1I**), confirming the method’s specificity for RNA-associated proteins.

### RTPP Analysis and validation of RNA-regulated proteins

To further validate the specificity of the identified RRPs, we meticulously compiled a comprehensive list of 5413 human RBPs annotated through prior RBP profiling methodologies^9–12,14,37–46^. RTPP demonstrated an RNA-binding specificity of 87%, surpassing the 77.3% specificity reported for R-DeeP (**Figure 2A, S2A**). These RRPs were annotated to interact with diverse RNA classes—including rRNA, tRNA, snRNA, and dsRNA—reflecting the method’s broad detection capability (**Figure 2B**). Interestingly, RRPs binding to rRNA exhibited significantly larger Δ*T*_m_ shifts compared to those interacting with other RNA types, suggesting stronger structural dependencies or more stable interactions with rRNA (**Figure 2C**).

**Figure 2.**
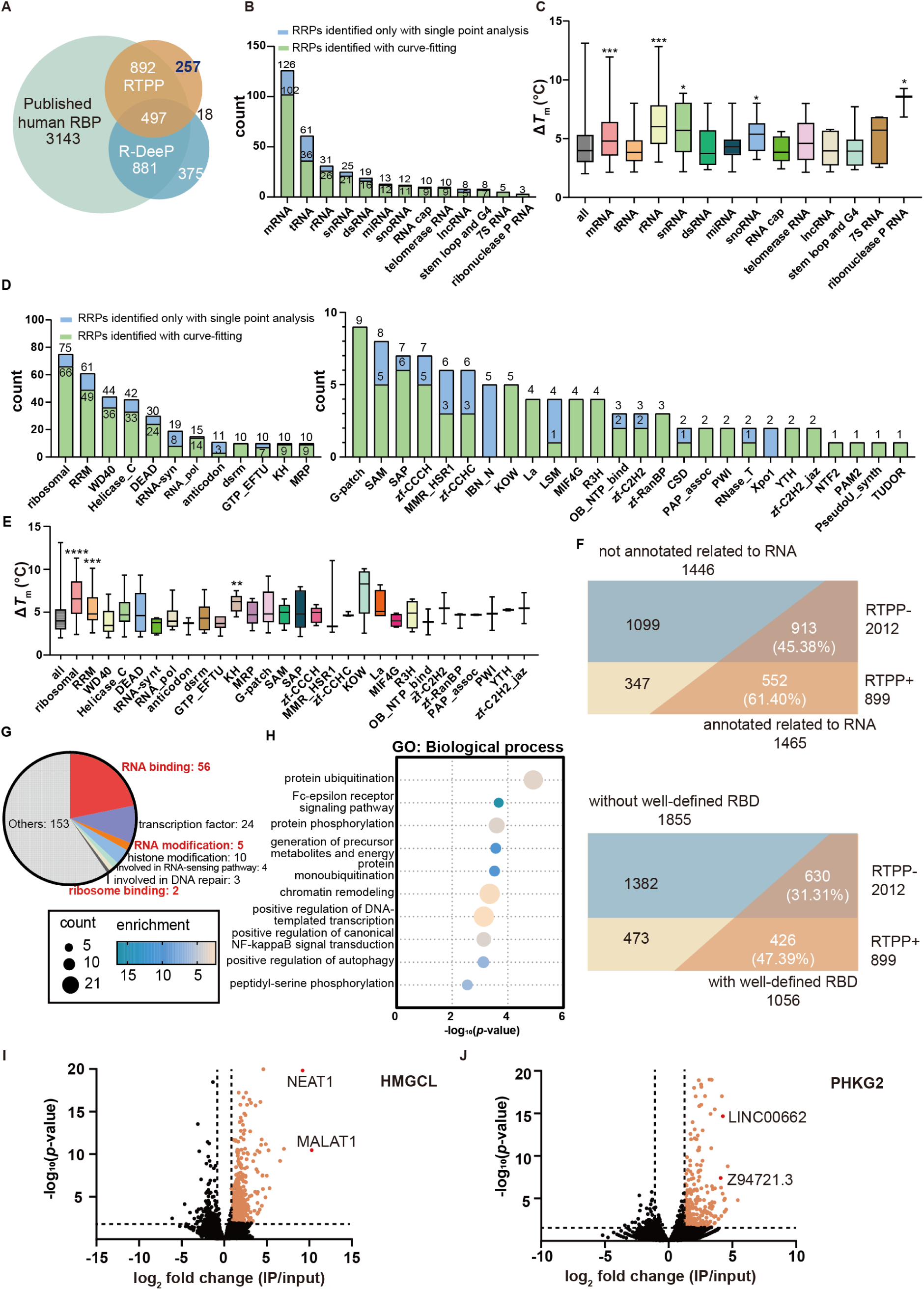
Analysis and validation of RNA-regulated proteins identified by RTPP. (A) Venn diagram showing the overlap of RNA-related proteins identified with traditional methods, R-DeeP, and RTPP. (B) Subclassification of RRPs with RNA binding species profile. (C) Comparison of Ä*T*_m_ values for RRPs bound to various types of RNA against. (D) Subclassification of RRPs with RNA binding domain profile. Terms with proteins less than 10 were separately shown. In (B) and (C), RRPs identified with shifted melting curve are shown in green, and those identified only by single temperature point comparison are shown in blue. (E) Comparison of Δ*T*_m_ values for RRPs containing typical RBDs. In (C) and (E), Error bars represent the mean ± s.d. \*\*\*\**p* < 0.0001, \*\*\**p* < 0.001, \*\**p* < 0.01, \**p* < 0.05 by student’s t-test against all RRPs. (F) Division of RBPs identified in crosslinking-based methods by whether identified with RTPP. The functionality of RBPs identified is defined by annotation in GO and having well-defined RBDs separately. In each case, percentage of functional RBPs in RTPP-identified group or RTPP-unidentified group were separately calculated. (G) Annotated evidence of RTPP-identified RBP orphans associated with various RNA-related functions. (H) GO-Biological Process analysis of RTPP-identified RBP orphans. (I) RIP-seq analysis of the RNA clients of HMGCL. (J) RIP-seq analysis of the RNA clients of PHKG2. In (I) and (J), significantly enriched RNAs are labeled in orange.

Among the RRPs, 517 contained classical RNA-binding domains (RBDs), with ribosomal and RNA recognition motifs (RRMs) being highly enriched (**Figure 2D**). Proteins with these domains showed larger Δ*T*_m_ shifts, potentially reflecting higher RNA-binding affinity (**Figure 2E**). Consistent with prior datasets, a notable fraction of mapped RRPs lack canonical RBDs. We also compared our results to crosslinking-based RBP catalogs from HEK293T cells^10,13,15,40^, finding that RTPP covered 899 previously reported RBPs in the same cellular context (**Figure S2B**). Crosslinking-detected RBPs not identified by RTPP tended to lack annotated RBDs or prior RNA-binding annotations, suggesting they may interact with RNA only transiently or non-specifically (**Figure 2F**).

RTPP uniquely identified 257 RRPs not previously captured by other methods (**Figure 2A**). Of these, 104 have published evidence supporting RNA association or roles in RNA-related processes, indicating they were likely missed due to limitations such as low UV-crosslinking efficiency (**Figure 2G**). Examples include HSF1, which binds a heat shock-responsive lncRNA^47^, thereby targeting stress genes, and the influenza NS1A-binding protein^48^, involved in RNA splicing and transport (**Figure S2C**). GO analysis revealed enrichment in terms related to ubiquitination and phosphorylation, hinting at novel RNA-dependent post-translational regulatory mechanisms (**Figure 2H**).

For experimental validation, we selected two novel RRPs without prior RNA-binding annotations: the mitochondrial enzyme HMGCL and the kinase subunit PHKG2 (**Figure S2D**). Western blot-based RTPP confirmed RNase-dependent thermal destabilization for both (**Figure S2E**). Orthogonal validation via UV-crosslinking and OOPS^15^ successfully recovered both proteins, even without crosslinking, confirming their RNA-binding activity (**Figure S2F**). RNA immunoprecipitation followed by sequencing (RIP-seq)^49^ further identified specific RNA interactors: HMGCL preferentially bound the lncRNAs MALAT1 and NEAT1, which have been reported in mitochondrial contexts^50^ (**Figure 2I**), while PHKG2 enriched the oncogenic lncRNAs LINC00662 and Z94721.1^51,52^ (**Figure 2J, Table S2**). These results validate RTPP as a robust method for discovering functional RNA-binding proteins.

### Long non-coding RNA CASC15 promotes SGK3 kinase activity by facilitating its PtdIns(3)P binding and endosomal recruitment

SGK3, a novel RNA-binding protein identified through RTPP (**Figure 3A**), is a Ser/Thr kinase structurally and functionally related to AKT^53^. It shares similar substrate specificity as AKT and has been implicated in tumorigenesis and immune responses^54–56^. A distinguishing feature of SGK3 is its N-terminal phox homology (PX) domain, which confers high-affinity binding to PtdIns(3)P—a lipid produced by class III PI3K (hVPS34) on endosomes in response to growth factors such as Insulin-Like Growth Factor-1 (IGF-1)^57,58^. The kinase activity of SGK3 depends on its endosomal localization, where it undergoes phosphorylation by PDK1 or mTORC2 and subsequently phosphorylates downstream substrates^59^. Western blot-based RTPP confirmed RNase treatment decreases the thermal stability of SGK3 (**Figure S3A**).

**Figure 3.**
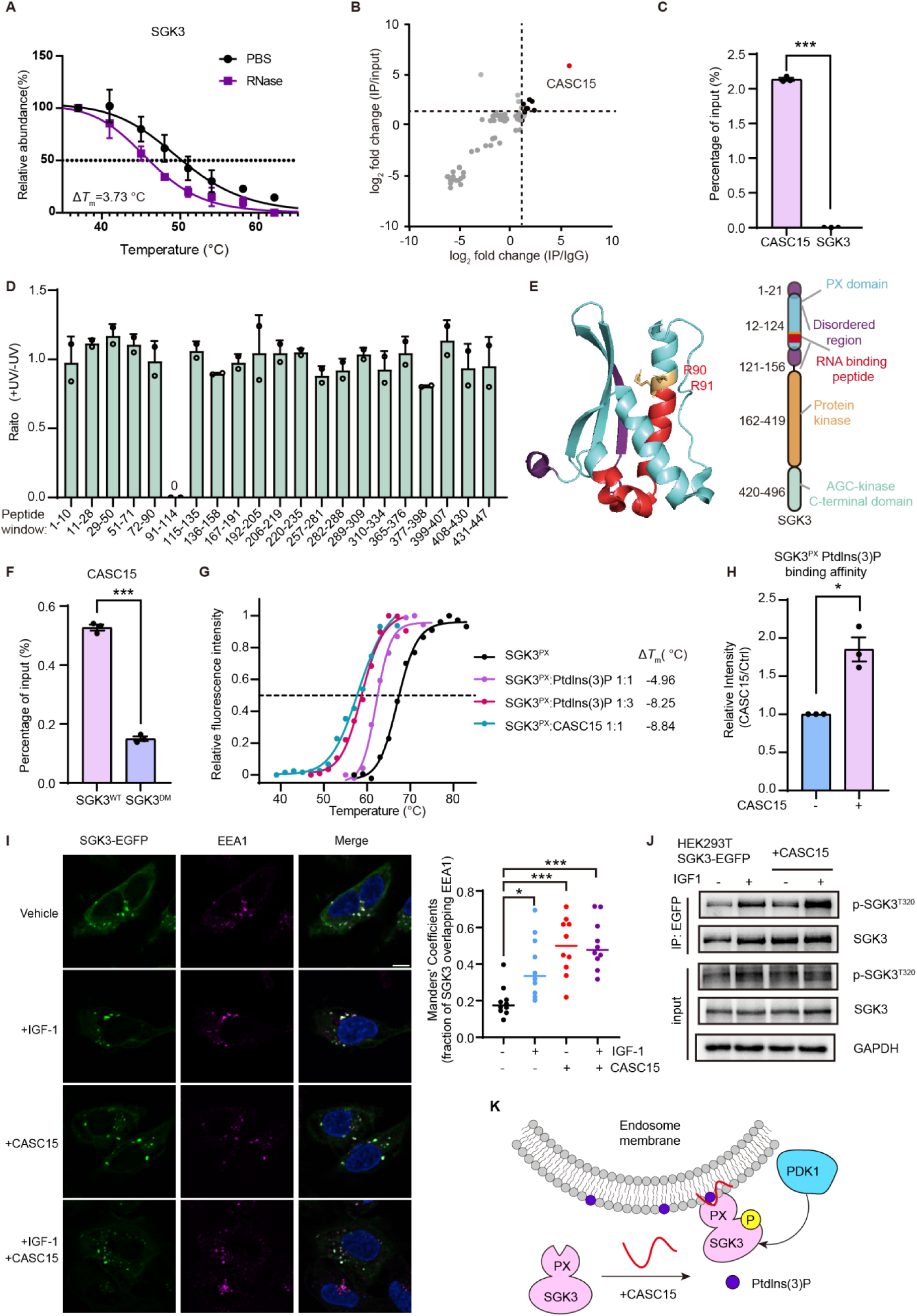
LncRNA CASC15 promotes SGK3 kinase activity by facilitating its PtdIns(3)P binding and endosomal recruitment. (A) Thermal melting curve of SGK3 in RTPP profiling. (B) RIP-seq analysis of SGK3 to identify its RNA clients. (C) Validation of CASC15 as a SGK3-associated RNA by RIP-qPCR analysis. The SGK3 RNA serves as a negative control. (D) Quantitative ratios (+UV vs. -UV) for all identified peptides from SGK3 in the RBR-ID assay. (E) Structure and domain information of SGK3. The potential RNA binding region is highlighted in red. (F) RIP-qPCR analysis of binding extent of wild-type SGK3 and double-mutant to CASC15. (G) Effect of Ptdlns(3)P and CASC15 on the thermal stability of purified PX domain. The purified PX domain was incubated with Ptdlns(3)P or CASC15 at indicated molecular ratios. The normalized *T*_m_ values were determined using a SYPRO Orange dye. (H) Quantification of the lipid dot blot assay. 10 nM SGK3 PX domain were incubated with 10 nM CASC15 for 1 hour, followed by UV crosslinking and incubation with nitrocellulose membrane pre-coated with 100 pmol Ptdlns(3)P. The dot blot images were presented in **Figure S3F**. Error bars represent the mean ± s.d. (A, C, D, F, H). \**p*<0.05, \*\**p*<0.01, \*\*\**p*<0.001 by two-sided student’s t-test (C, F, and H) (I) Immunofluorescence of SGK3-EGFP and the endosome marker EEA1 in U2OS cells upon CASC15 overexpression and IGF-1 stimulation. The fraction of SGK3-EGFP overlapping with endosomes were quantified from 10 cells for each experimental condition. \**p*<0.05, \*\*\**p*<0.001 by ordinary one-way ANOVA. Scale bars, 10 μm. (J) T320 phosphorylation of SGK3-EGFP in HEK293T cells upon CASC15 overexpression and IGF-1 stimulation. SGK3-EGFP was immunoprecipitated using anti-EGFP agarose resins and blotted against indicated antibodies. (K) Proposed working model of CASC15 facilitating the recruitment of SGK3 to endosome membrane through binding at the PX domain.

To identify RNAs that interact with SGK3 under basal and IGF-1-stimulated conditions, we performed RIP-seq in MCF-7 cells. This revealed the tumor-associated lncRNA CASC15 as the most significantly enriched SGK3-binding RNA (**Figure 3B**). IGF-1 stimulation also altered the RNA-binding profile of SGK3 (**Figure S3B, Table S2**), suggesting a potential link between its endosomal recruitment and RNA binding. Although CASC15 is known to be involved in various cancers and represents a promising therapeutic target, its protein partners and regulatory mechanisms remain poorly understood^60,61^. To validate SGK3 as a novel binding partner of CASC15, we performed RIP followed by quantitative RT-PCR (**Figure 3C**), which confirmed the interaction between SGK3 and CASC15. Given the shared relevance of SGK3 and CASC15 in tumorigenesis, their interaction may reveal new therapeutic avenues.

To map the RNA-binding region on SGK3, we used the RBR-ID assay^39^, which relies on UV crosslinking of RNA to bound peptides, leading to a reduction in abundance of the corresponding unmodified peptides (**Figure S3C**). UV crosslinking specifically decreased the abundance of peptides spanning Arg91 to Arg114, while other peptides remained unaffected, indicating that this region constitutes the RNA-binding site (**Figure 3D**). This segment lies within the PX domain and is flanked by intrinsically disordered regions (IDRs) (**Figure 3E**), consistent with the frequent enrichment of IDRs near RNA-binding sites. Given the critical role of arginine residues in RNA–protein interactions, we mutated Arg90 and Arg91 to alanine and RIP–qPCR demonstrated that this mutation significantly impaired SGK3 binding to CASC15 (**Figure 3F**).

Because SGK3 binds both RNA and PtdIns(3)P via its PX domain, we asked whether CASC15 influences lipid binding. We purified the SGK3 PX domain (**Figure S3D**) and performed a SYPRO Orange-based thermal shift assay in the presence of purified CASC15 or PtdIns(3)P. Both treatments reduced the thermal stability of the PX domain (**Figure 3G**), suggesting similar binding modes. CASC15 had a greater destabilizing effect than PtdIns(3)P, implying potentially stronger binding affinity. Using a dot blot assay with lipid-coated nitrocellulose membranes (**Figure S3E**), we confirmed that the PX domain specifically binds PtdIns(3)P, but not PtdIns(4)P (**Figure S3F**). Notably, pre-incubating the PX domain with CASC15 enhanced its binding to PtdIns(3)P, indicating that CASC15 facilitates this interaction (**Figure 3H**).

To assess this effect in a cellular context, we generated U2OS cells stably expressing EGFP-tagged wild-type SGK3 (SGK3^WT^) or the RNA-binding-deficient double mutant (SGK3^DM^). SGK3^WT^ localized predominantly to endosomes, whereas SGK3^DM^ was largely cytosolic (**Figure S3G**). Consistent with our i*n vitro* data, overexpression of CASC15 promoted the endosomal localization of SGK3 (**Figure 3I**). Since endosomal recruitment is essential for SGK3 activation, we immunoprecipitated SGK3-EGFP and found that CASC15 overexpression increased phosphorylation at Threonine 320—a marker of SGK3 activity—under both basal and IGF-1-stimulated conditions (**Figure 3J**). In summary, we demonstrate that the lncRNA CASC15 binds to the PX domain of SGK3, enhances its interaction with PtdIns(3)P, promotes its recruitment to endosomes, and thereby potentiates its kinase activity **(Figure 3K)**.

### An organ-wide atlas of RNA-regulated proteins in mice

Most existing RBP profiling methods rely on UV crosslinking, which cannot be readily applied to intact tissues due to poor light penetration. Consequently, the global landscape of RNA-binding proteins across mammalian organs remains largely unexplored. Given that TPP is compatible with tissue samples^26^, we applied RTPP to five adult mouse organs—brain, lung, liver, kidney, and spleen—to systematically map RRPs under physiological conditions (**Figure 4A**). Using DIA-based LC-MS/MS, we quantified 6,641 proteins with Δ*T*_m_ values across all organs (**Figure S4A, B**). Applying stringent thresholds (Δ*T*_m_ > 2°C, *p* < 0.05), we identified high-confidence RRPomes ranging from 149 proteins in the liver to 735 in the lung, with 1,372 RRPs identified in total (**Figure 4B, S4C, Table S3**). Notably, only 447 of these overlapped with RRPs identified in HEK293T cells, highlighting the importance of direct tissue profiling (**Figure S4D**).

**Figure 4.**
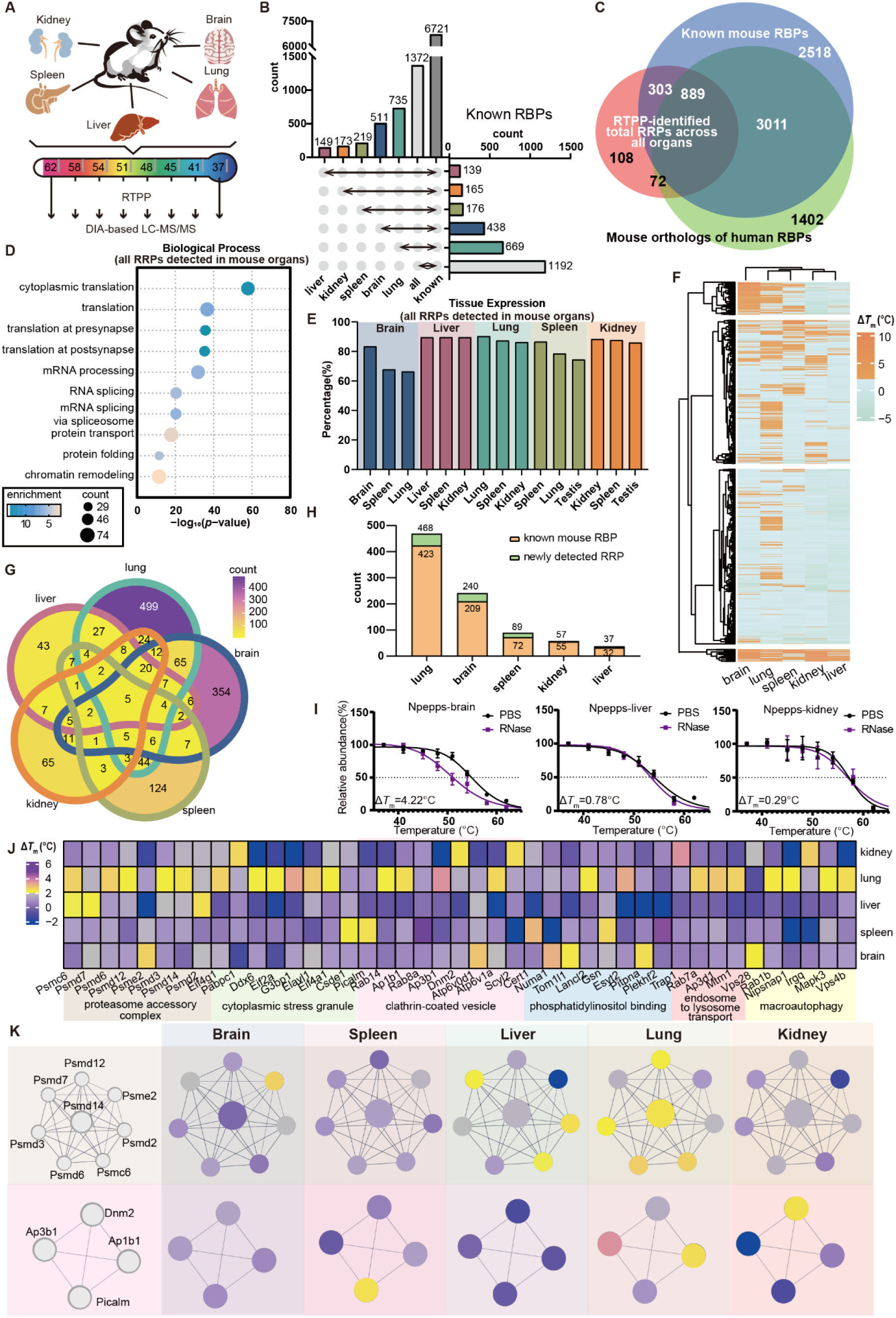
Atlas of RNA-regulated proteins across mouse organs. (A) Schematic of RTPP profiling in five different mouse organs: brain, kidney, liver, lung and spleen. (B) Numbers of RRPs identified in each organ and numbers of published mouse RBPs in each organ. (C) Venn diagram showing the overlap of all RRPs identified in mouse organ RTPP assays, all mouse RBPs published, and mouse orthologs of all published human RBPs. (D) GO-Biological Process analysis of all RRPs identified in organs. (E) GO-Tissue Expression analysis of RRPs in different organs. (F) Heatmap showing hierarchical clustering of Δ*T*_m_ values for RRPs identified in five organs. (G) Venn diagram showing the overlap among RRPs identified in organs. The color of each region indicates the number of RRPs within the region. (H) Numbers of organ-specific RRPs and numbers of published mouse RBPs within them. (I) Thermal melting curves of a representative brain-specific RRP, NPEPPS, in distinct organs. Error bars represent the mean ± s.d. (J) Heatmap of selected representative organ-specific RRPs involved in various biological processes and complexes. (K) Interaction map of two representative clusters of proteins shown in (J). The color of each node indicates its Δ*T*_m_ value in the corresponding organ.

The majority of mouse RRPs (86.9%) were previously annotated as RBPs^12,62–66^, and an additional 72 were orthologs of known human RBPs (**Figure 4C**). GO analysis confirmed strong enrichment of RNA-related processes such as mRNA processing, splicing, and translation (**Figure 4D**). Shared RRPs across multiple organs were enriched in housekeeping RNA-binding functions, including ribosomal and spliceosomal components (**Figure S4E**). As observed in cell culture, crosslinking-detected RBPs not identified by RTPP tended to lack prior RNA-binding annotations, suggesting transient or indirect interactions (**Figure S4F**). Newly discovered RRPs showed enrichment in innate immunity, mitophagy, and intracellular transport, pointing to expanded roles for RNA *in vivo* (**Figure S4G**).

The RTPP-derived mouse RRPomes correctly highlighted proteins overrepresented in specific organs, demonstrating high tissue specificity (**Figure 4E**). We also observed significant organ-to-organ variation in Δ*T*_m_ values, with many RRPs showing RNA dependence in only a single tissue (**Figure 4F**). This points to an unexpectedly high degree of organ-specific heterogeneity in RNA binding. Indeed, while only five RRPs were common to all five organs, 1,085 were identified in just one (**Figure 4G**). These organ-specific RRPs may include proteins uniquely expressed in that tissue, or ubiquitously expressed proteins whose thermal stability becomes RNA-dependent in an organ-specific manner. The lung contained the highest number of organ-specific RRPs (468), followed by the brain (240), spleen (89), kidney (57), and liver (37), the majority of which are previously annotated RBPs (**Figure 4H**). For example, puromycin-sensitive aminopeptidase (NPEPPS), which digests tau in the mouse brain, exhibited RNase-sensitive thermal stability exclusively in brain tissue and not in other organs (**Figure 4I**)^67^. These organ-specific RRPs were enriched for functions related to RNA processing **(Figure S4H)**, and were also enriched for multiple protein complexes not typically associated with RNA, such as the proteasome accessory complex and clathrin-coated vesicles (**Figure 4J**). Intriguingly, many of these complexes exhibited distinct RNA-associated components depending on the organ (**Figure 4K**), further emphasizing the tissue-dependent nature of RNA-protein interactions.

### Identification of Alzheimer’s disease-relevant RNA-regulated proteins in mouse hippocampus

Building on the tissue applicability of RTPP, we next sought to characterize aberrant RNA–protein interactions in disease. Alzheimer’s disease (AD) involves dysregulated RNA metabolism and impaired RNA-binding protein function^68^. Although proteomic studies have revealed expression changes in AD brains^69,70^, alterations in protein thermal stability and RNA binding during disease progression remain poorly understood. To address this, we performed RTPP on hippocampal tissue from 5xFAD transgenic AD mice^71^ and wild-type controls (**Figure 5A**). DIA-based analysis quantified 4,540 proteins in both normal and AD hippocampi (**Figure S5A**). Using standard thresholds (Δ*T*_m_ > 2°C, *p* < 0.05), we identified 690 RRPs in the normal hippocampus and 3,418 in the AD hippocampus, indicating a marked global increase in RNA–protein interactions during AD progression (**Figure S5B, Table S4**). These RRPs showed strong overlap with established RBP datasets, supporting their RNA-binding specificity (**Figure S5C, D**).

**Figure 5.**
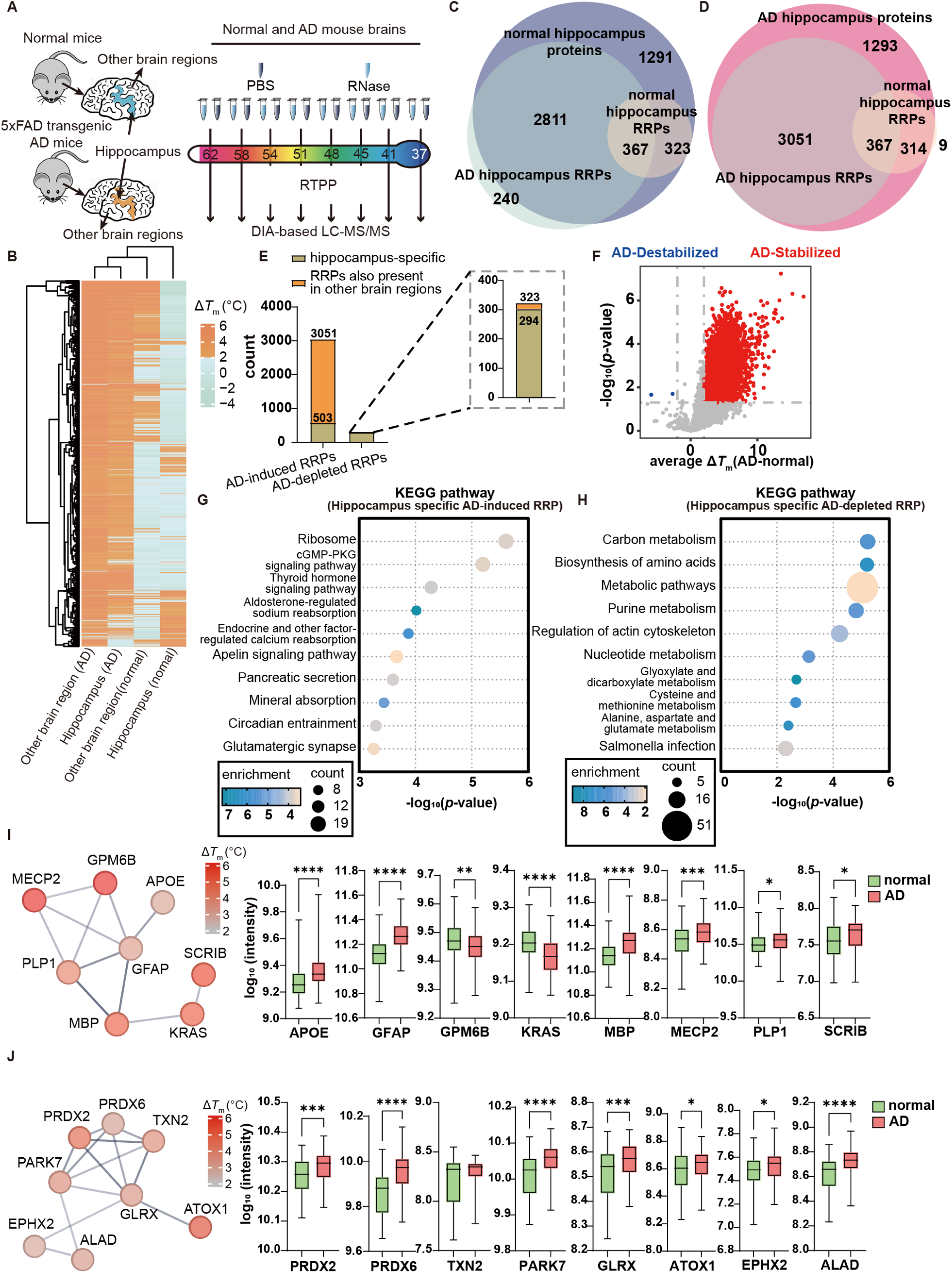
Identification of Alzheimer’s disease-relevant RNA-regulated proteins in mouse hippocampus. (A) Schematic of hippocampus-specific RTPP profiling in normal and 5xFAD mouse brains. (B) Heatmap showing hierarchical clustering of RRPs identified in the hippocampus and other brain regions of normal and AD mice. (C) Venn diagram showing the overlap of RRPs identified in the normal hippocampus, RRPs identified in the AD hippocampus, and total proteins quantified in the normal hippocampus. (D) Venn diagram showing the overlap of RRPs identified in the normal hippocampus, RRPs identified in the AD hippocampus, and total proteins quantified in the AD hippocampus. (E) Hippocampus specificity of AD-induced and AD-depleted RRPs. (F) Volcano plot of total quantified proteins with Δ*T*_m_ values in the normal and AD hippocampus. Proteins with significantly altered thermal stability are colored in red and blue. (G) KEGG analysis of hippocampus-specific AD-induced RRPs (H) KEGG analysis of hippocampus-specific AD-depleted RRPs (I) Representative AD-induced RRPs with change in protein intensity between AD patients and normal people in Banner cohort, enriched in synapse organization and myelin sheath. (J) Representative AD-depleted RRPs with change in protein intensity between AD patients and normal people as control in Banner cohort, mainly enriched in oxidative stress response. In (I) and (J), colors of the nodes indicate the Δ*T*_m_ value of the protein. \*\*\*\**p* < 0.0001, \*\*\**p* < 0.001, \*\**p* < 0.01, \**p* < 0.05 by Wilcoxon rank-sum test (I, J).

We observed pronounced differences in Δ*T*_m_ between normal and AD conditions (**Figure 5B**). Beyond the 367 RRPs common to both, we identified 3,051 AD-induced RRPs (increased RNA dependence in AD) and 323 AD-depleted RRPs (lost RNA dependence in AD). Among AD-induced RRPs, 2,811 showed RNase-dependent destabilization only in AD tissue, while 240 were uniquely quantified in AD samples (**Figure 5C**). Conversely, 314 AD-depleted RRPs lost RNA-dependent stability specifically in AD, and 9 were absent in AD hippocampi (**Figure 5D**). To define hippocampus-specific disease-associated RRPs, we compared hippocampal data with other brain regions. This revealed 503 hippocampus-specific AD-induced and 294 hippocampus-specific AD-depleted RRPs (**Figure 5E**). Additionally, even without RNase treatment, many proteins exhibited altered thermal stability in AD hippocampi, consistent with reported proteome-wide instability in AD^72^ (**Figure 5F**).

Functional enrichment analysis showed that hippocampus-specific AD-induced RRPs were associated with ribosomal function, cGMP-PKG signaling, and thyroid hormone pathways (**Figure 5G**). In contrast, AD-depleted RRPs were enriched in metabolic processes including carbon metabolism, amino acid biosynthesis, and purine metabolism (**Figure 5H**)—pathways known to be disrupted in AD.

To assess human relevance, we analyzed proteomic data from the Banner cohort, correlating RRP changes with CERAD and Braak neuropathological scores^73,74^. We found that a significant proportion of AD-dependent RRPs are dysregulated in AD patients (**Figure S5F**), and components of molecular networks are co-regulated during AD progression. For instance, synaptic components (enriched among AD-induced RRPs) and oxidative stress regulators (enriched among AD-depleted RRPs) showed correlation with disease progression (**Figure 5I, J, S5F–H**). Thus, by leveraging its compatibility with intact tissue, RTPP enables the identification of disease-associated RNA–protein interactions and uncovers potential mechanisms underlying AD pathogenesis.

### Revealing subcellular heterogeneity of RNA regulation via spatiotemporally resolved RTPP

Compartmentalization of eukaryotic cells and the dynamic distribution of macromolecules, such as RNA and proteins, across these compartments are crucial for cellular function^75^. Many RBPs localize in multiple organelles but may exhibit distinct interactions with RNA; however, current RBP profiling methods lack spatiotemporal resolution^45^. After addressing the heterogeneity of RNA binding at tissue- and disease-dependent levels, we aimed to investigate spatially restricted RNA–protein interactions and identify organelle-specific RRPs. To accomplish this, we integrated RTPP with TurboID-based proximity biotinylation, a non-toxic method to label subcellular proteomes with nanometer spatial resolution^76^(**Figure 6A**). HEK293T cells were transfected with TurboID fused to nuclear localization signal (NLS) and nuclear export signal (NES) to label nuclear and cytosolic proteins, respectively. Their labeling specificity was confirmed through fluorescent imaging (**Figure S6A**). Following RNase treatment and thermal denaturation, biotinylated proteins were isolated via streptavidin pull-down, with validation confirming that this enrichment did not distort thermal stability measurements (**Figures S6B, C**).

**Figure 6.**
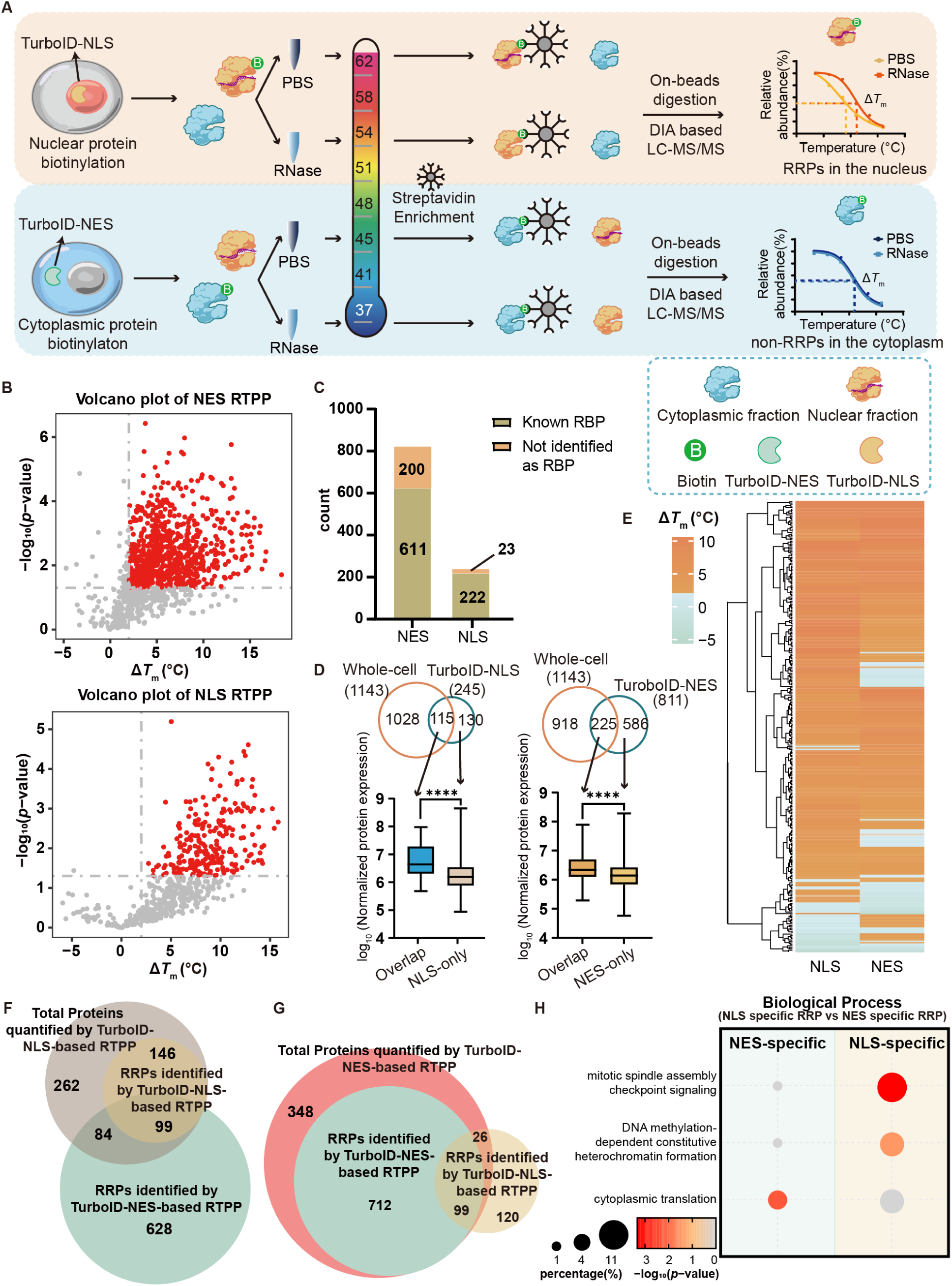
Spatiotemporally resolved profiling of subcellular RNA-regulated proteins via proximity labeling-coupled RTPP. (A) Schematic of PL-RTPP for mapping subcellular RRPs. HEK293T cells expressing TurboID-NLS or TurboID-NES were labeled with biotin, then the cell lysates were treated separately with PBS or RNase, divided into aliquots and heated to the indicated temperatures. Supernatants were subjected to streptavidin enrichment, on-beads trypsin digestion, and DIA-based LC-MS/MS analysis. (B) Volcano plot of proteins quantified with Δ*T*_m_ by TurboID-NES-based and TurboID-NLS-based RTPP. Proteins with Δ*T*_m_ > 2°C and *p*-value < 0.05 were annotated as RRPs and labeled in red. Statistical analysis was performed using a two-sided student’s t-test. (C) Number of known RBPs identified as nuclear or cytosolic RRPs identified by PL-RTPP. (D) Overlap of RRPs identified by whole-cell RTPP and subcellular RRPs identified by PL-RTPP. Box plots compare the protein abundance between RRPs detected in both datasets and RRPs detected only by PL-RTPP. \*\*\*\**p*<0.0001 by a two-sided Wilcoxon rank-sum test (E) Heatmap showing hierarchical clustering of RRPs identified by TurboID-NES-based RTPP and TurboID-NLS-based RTPP. (F) Overlap between RRPs identified by TurboID-NES-based RTPP, RRPs identified by TurboID-NLS-based RTPP, and total proteins quantified by TurboID-NES-based RTPP. (G) Overlap between RRPs identified by TurboID-NES-based RTPP, RRPs identified by TurboID-NLS-based RTPP, and total proteins quantified by TurboID-NLS-based RTPP. (H) Representative GO terms specifically enriched in NES/NLS-specific RRPs. The size of each bubble represents the number of RRPs in the term, and the color gradient indicates the percent of RRPs.

Quantitative proteomics identified 245 nuclear and 811 cytosolic RRPs exhibiting significant RNase-dependent thermal destabilization (**Figure 6B and Table S5**). The vast majority (91% nuclear, 75% cytosolic) were previously known RBPs, and GO analysis confirmed strong enrichment of RNA-related functions (**Figure 6C and S6D**). GO subcellular compartment analysis reveals a specific enrichment of terms related to the nucleus and cytosol, revealing that some proteins are localized in both compartments (**Figure S6E**). We found that only 115 nuclear RRPs (47%) and 225 cytosolic RRPs (28%) were identified by whole-cell RTPP; those RRPs not detected by whole-cell RTPP tended to be lower-abundance proteins (**Figure 6D**). We surmise that by focusing on a single subcellular compartment, PL-RTPP might probe more deeply than global RTPP profiling and identify lower-abundance RBPs more effectively.

Crucially, many proteins displayed compartment-specific RNA-binding behavior. While 99 RRPs were shared, 84 were RNA-bound only in the cytosol and 26 only in the nucleus (**Figures 6E–G**). These spatially restricted RRPs were linked to compartment-specific processes: nuclear RRPs to chromatin organization and mitotic spindle assembly, and cytosolic RRPs to translation (**Figure 6H**). Furthermore, 81 compartment-specific RRPs were missed entirely by global profiling (**Figure S6F**), likely obscured by non-binding isoforms elsewhere. Thus, PL-RTPP offers a valuable approach to resolve subcellular RNA-binding heterogeneities and discover localized RBPs.

### Development of cell surface RTPP to identify proteins regulated by cell surface RNA

While RNA-protein interactions within the cell are well characterized, the reciprocal regulatory relationships between RNAs and proteins on the cell surface remain largely unexplored. Recent studies have identified cell surface RNAs, such as glycoRNAs, which play roles in intercellular communication and immune responses^77–80^. Notably, glycoRNAs form nanoclusters with cell surface RNA-binding proteins (csRBPs)^78^. However, csRBPs identified to date have been inferred primarily by cross-referencing known cell surface protein and RBP datasets—an approach that lacks direct evidence of RNA binding or functional regulation. Furthermore, current RBP profiling methods are unsuitable for csRBPs due to several challenges: csRNAs are mostly small, non-polyadenylated RNAs, precluding oligo-dT pulldown; their low abundance hinders efficient metabolic labeling; and glycosylation modifications on both RNA and proteins interfere with organic-aqueous phase separation.

To directly map proteins regulated by csRNAs, we developed a cell surface variant of RTPP (csRTPP). Inspired by live-cell RNase A assays used in glycoRNAs studies which selectively cleave extracellular RNAs while leaving intracellular RNAs intact—we applied RNase treatment directly to intact cells rather than lysates. This approach allows us to link molecular changes to phenotypic observations from live-cell RNase experiments (**Figure 7A**). To quantify thermal stability shifts in cell surface proteins, we lysed RNase A-treated HEK293T cells in PBS containing 0.4% NP-40, adapting a protocol previously established for plasma membrane protein TPP^28^. Western blot analysis of DDX21 and HNRNPU—two validated csRBPs known to form glycoRNA nanoclusters—revealed a significant reduction in thermal stability following live-cell RNase treatment (**Figure 7B**). In contrast, GAPDH stability remained unchanged, confirming the specificity of csRTPP.

**Figure 7.**
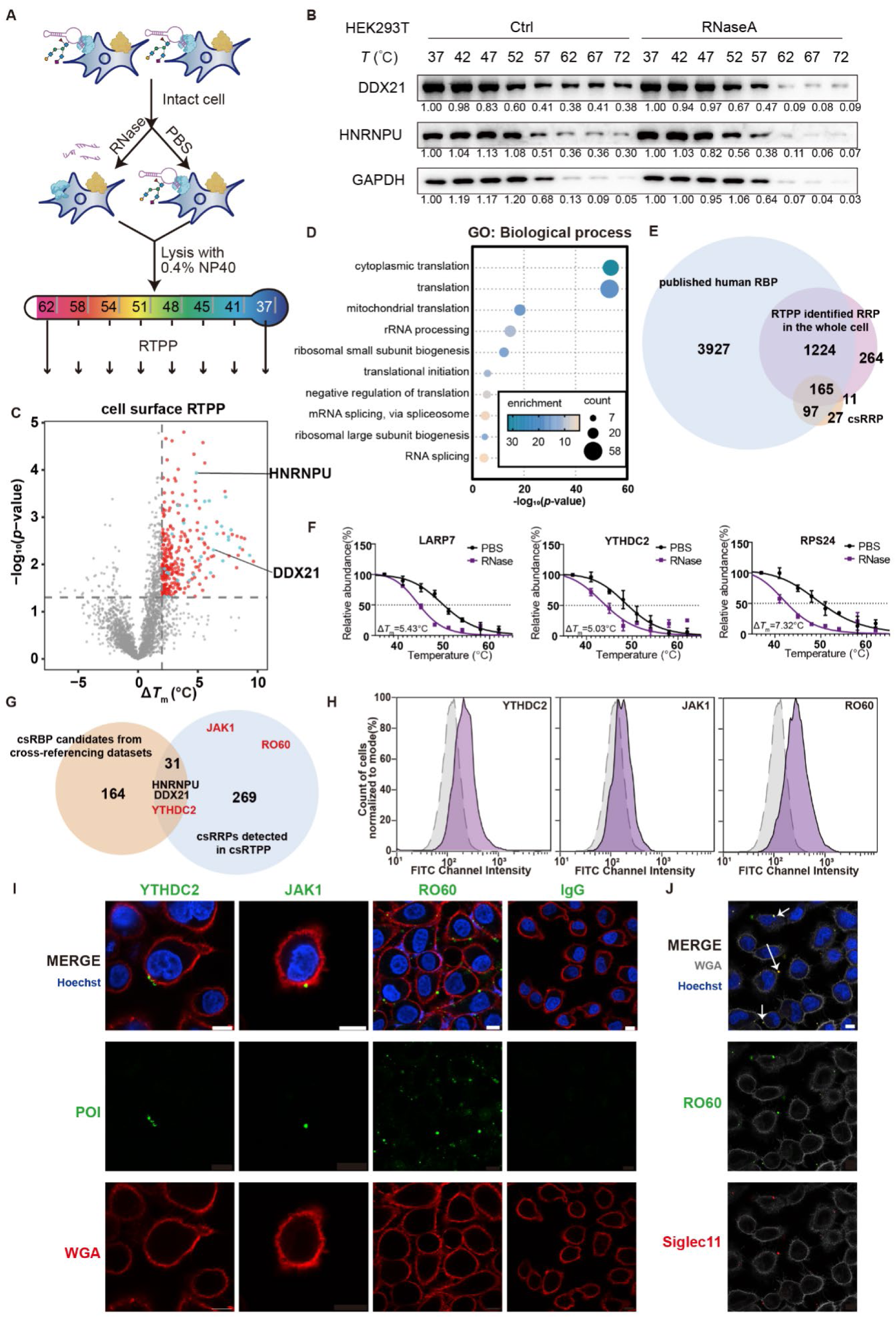
Development of cell surface RTPP reveals nanoscale clusters of RNA-protein complexes on living cells. (A) Schematic of cell surface RTPP (csRTPP) profiling on intact cells. Live cells were treated with RNase to cleave cell surface RNAs, followed by lysis with 0.4% NP-40 and thermal proteome profiling. (B) Western blot analysis of two previously reported csRBPs and GAPDH following the csRTPP assay. (C) Volcano plot of proteins quantified with Δ*T*_m_ in HEK293T cells using csRTPP. Proteins with Δ*T*_m_ > 2°C and *p*-value < 0.05 were annotated as csRRPs and labeled in red. Among these, csRRPs also identified as csRBPs in cross-reference datasets are labeled in blue. Statistical analysis was performed using a two-sided student’s t-test. (D) GO Biological Process analysis of csRRPs identified in HEK293T cells. (E) Venn diagram showing the overlap of csRRPs in HEK293T cells, RRPs identified in HEK293T cells with the whole-cell RTPP profiling, and all previously reported human RBPs. (F) Thermal melting curves of three representative csRRPs involved in ribosome function or RNA modification recognition. Error bars represent the mean ± s.d. (G) Comparison of csRRPs identified by csRTPP with proteins previously annotated as csRBPs from cross-referencing datasets. (H) FACS histograms of three candidate csRRPs compared to the control group. (I) Confocal microscopy images of candidate csRRPs co-stained with WGA and Hoechst33342 on live cells. Scale bar, 10ìm. (J) Confocal microscopy images of RO60, co-stained with siglec-11, along with WGA and Hoechst33342 on live cells. Scale bar, 10ìm.

DIA-based proteomics identified 300 proteins with significant thermal destabilization (Δ*T*_m_ > 2°C, *p* < 0.05) (**Figure 7C, S7A and Table S6**). GO analysis indicated that these cell surface RNA-regulated proteins (csRRPs) were highly enriched for RNA-related functions (**Figure 7D and S7B**). Comparison with established RBP datasets demonstrated an RNA-binding specificity of 87.3% (**Figure 7E**). Consistent with prior cross-referencing studies, most csRRPs were classical intracellular RBPs, including ribosomal proteins and RNA modification readers (**Figure 7F, S7C**). These proteins possess well-defined RNA-binding domains that are typically absent in plasma membrane proteins (**Figure S7D**). Notably, 27 csRRPs had not been previously annotated as RBPs or detected by whole-cell RTPP, and included several mitochondrial proteins (**Figure S7E**). These may represent proteins with only a small fraction localized to the cell surface in association with RNA. Independent cell surface proximity labeling studies also report an overrepresentation of mitochondrial proteins^81^, suggesting the existence of uncharacterized pathways that deliver intracellular and mitochondrial proteins to the cell surface.

### csRRPs form nanoscale clusters with glycoRNAs on the cell surface

A direct comparison with csRBPs identified through the data-crossing approach revealed only 31 overlapping proteins, including DDX21 and HNRNPU (**Figure 7G**). This limited overlap underscores the importance of direct mapping for identifying authentic csRBPs. The proteins uniquely identified by csRTPP include numerous RBPs known to interact with diverse RNA types, such as rRNA, snRNA, and tRNA (**Figure S7F**). For instance, csRTPP detected RO60, an RNA-binding protein that interacts with intracellular Y RNAs and rRNAs—both of which are also present on the cell surface^82^. We also identified the 3’-5’ RNA helicase YTHDC2, a recognized reader of m6A modification, as a csRRP, suggesting potential epitranscriptomic regulation on cell surface RNAs. Notably, csRTPP revealed that the tyrosine-protein kinase JAK1—a critical kinase partner for several interferon receptors traditionally considered to be intracellular—also localizes to the cell surface and appears to interact with cell surface RNAs.

We validated the cell surface localization of RO60, YTHDC2, and JAK1 by incubating live cells with validated antibodies against each protein, followed by flow cytometry analysis (FACS). All three proteins exhibited surface expression, with RO60 showing the highest signal contrast (**Figure 7H**). After live-cell antibody incubation, cells were fixed and co-stained with fluorophore-conjugated Wheat Germ Agglutinin (WGA). Consistent with previous observations for HNRNPU^78^, all three proteins were detected on the cell surface and formed nanoscale clusters (**Figure 7I**). Z-stack imaging revealed distinct cluster morphologies: RO60 formed numerous small puncta, whereas JAK1 assembled into substantially larger structures (**Figure S7G**).

The classical binding partner of RO60, Y5 RNA, regulates its subcellular localization and is critical for cellular stress responses. Given that Y5 RNA is a prevalent glycoRNA on the cell surface, we asked whether RO60 co-localizes with Y5 RNA in this compartment. Using a fluorophore-conjugated hybridization probe targeting Y5 RNA, we found that Y5 RNA forms nanoscale clusters on the surface of live cells, which were abolished by RNase treatment (**Figure S7H**). Importantly, Y5 RNA signals consistently co-localized with or localized adjacent to RO60 puncta (**Figure S7I**). Co-staining with Siglec-11—a recombinant protein consisting of the Siglec-11 extracellular domain fused to a human Fc domain, which binds specifically to cell surface glycoRNA^83^—further demonstrated clear co-localization between RO60 and glycoRNA signals (**Figure 7J**). Together, these data support the hypothesis that Y5 RNA and RO60 may translocate and function together on the cell surface.

## DISCUSSION

The identification of novel RNA-associated proteins can provide insights into previously unknown RNA-dependent regulatory mechanisms. Over the past decade, numerous proteomic methods have been developed to discover RBPs, most of which rely on UV crosslinking and the enrichment of RNA-protein complexes^9–15,21^. While these methods have yielded valuable RBP datasets from mammalian cells, the specific mechanisms behind RNA-protein interactions remain largely unclear for many of the identified proteins. The sheer number of RBPs discovered also raises concerns that some may simply be incidentally crosslinked to RNA during UV treatment, potentially resulting in false positives that do not functionally interact with RNA. There is an urgent need for more refined datasets of RBPs that are more likely to be functionally engaged with RNA.

The RTPP technology we present here detects structural changes in proteins when RNA is depleted, directly reflecting the functional impact of RNA on its cognate protein partners. While RTPP does not necessarily identify direct RNA binding events, the majority of the RRPs identified by RTPP were previously annotated as such by UV-based methods. These RBPs are likely to be more functionally relevant to RNA compared to those not identified by RTPP, which may only interact with RNA sporadically or at very low stoichiometry. Additionally, RTPP identifies 257 RRPs that were missed by all prior RBP profiling approaches. Further literature analysis confirms that a significant portion of these proteins functionally associate with RNA, even if they do not directly bind to it. To explore this further, we performed RIP-seq on three RRPs—HMGCL, PHKG2 and SGK3—that were not previously known to interact with RNA. In particular, in-depth investigations reveal that SGK3 binds to the long non-coding RNA CASC15 at its PtdIns(3)P binding site, which promotes SGK3’s endosome recruitment and kinase activity. Beyond functional relevance, RTPP also outperforms other RBP profiling methods in terms of required starting materials and processing time. Furthermore, since the CETSA assay is widely used to evaluate drug-target engagement^25,26^, we believe that the Western blot-based RTPP can be easily adopted by most biochemical laboratories to validate whether an RBP binds RNA inside cells.

Another key feature that makes RTPP unique is its ability to be easily applied to tissue samples, a capability that most previous crosslinking-based methods cannot achieve. As a result, we provide a valuable atlas of RRPs across various mouse organs, significantly expanding the scope of RNA-protein interactions by identifying over 1,00 novel RNA-associated proteins. Notably, we observed a substantial number of tissue-specific RRPs. These proteins are expressed in multiple organs but only bind RNA in specific tissues, highlighting organ-dependent variations in RNA regulation. We further applied RTPP to profile RRPs in the hippocampus of AD mice, revealing AD-specific RNA-protein interactions within this brain region. Many of these AD-dependent RRPs are highly relevant to disease progression, suggesting that dysregulated RNA-protein interactions may contribute to AD pathology. These *in vivo* findings underscore the utility of RTPP in exploring the heterogeneity of RNA-protein interactions across tissues, brain regions, disease states, and beyond.

To further explore the heterogeneity of RNA-protein interactions at the subcellular level, we combine RTPP with proximity biotinylation for spatiotemporal analysis of RRPs. Our results demonstrate that PL-RTPP can identify compartment-specific RRPs, which are expressed in multiple cellular compartments but exhibit RNA-dependence in only one. We also developed csRTPP to detect RRPs associated with cell surface RNA via live-cell RNase treatment. Surprisingly, this approach revealed the cell surface localization of several intracellular RRPs. Imaging validation of three candidates—RO60, YTHDC2, and JAK1—showed that they form nanoscale clusters on the cell surface and colocalize with cognate glycoRNAs. The mechanisms underlying the formation and functional roles of these molecular clusters warrant further investigation. We anticipate that RTPP will be widely adopted to uncover functional RNA-binding events and elucidate their heterogeneity across diverse biological contexts.

### Limitations of the study

Since RTPP detects changes in protein structural stability upon RNA depletion, it may also capture indirect RNA-binding events. Although most of the RTPP-identified RRPs are known RBPs, our datasets likely include proteins that do not directly bind RNA but interact with other RBPs. While some of these indirect RRPs are functionally relevant to RNA, false positives may also be present. Additionally, we observed proteins that exhibit increased thermal stability upon RNase treatment. We hypothesize that some of these proteins may directly bind RNase and become stabilized, while others may show decreased thermal stability upon RNA binding.

The sensitivity of RTPP in detecting RNA dependence is influenced by RNA-binding stoichiometry. If only a small fraction of a protein is bound to RNA, structural changes may be obscured by the unbound population. Although proximity labeling helps distinguish RNA-binding behaviors across subcellular regions, it cannot resolve differences among certain proteoforms—such as post-translationally modified versus unmodified variants—that may exhibit distinct RNA-binding characteristics. This limitation currently falls outside the scope of RTPP alone. However, future integration of RTPP with complementary approaches, such as phosphoproteomics, may help address this challenge.

## RESOURCE AVAILABILITY

### Lead contact

Requests for further information and resources should be directed to and will be fulfilled by the lead contact, Wei Qin.

### Materials availability

All unique/stable reagents generated in this study are available from the lead contact with a completed materials transfer agreement.

### Data and code availability

Proteomic data have been deposited at iProx at IPX0014638000 and are publicly available as of the date of publication.

Original western blot images and microscopy data reported in this paper will be shared by the lead contact upon request.

Any additional information required to reanalyze the data reported in this paper is available from the lead contact upon request.

## ACKNOWLEDGMENTS

The authors would like to acknowledge Dr. Yuan Liu, College of Chemistry and Molecular Sciences, Peking University, for help with fitting the melting curves. We would like to thank the Image Core Facility, Technology Center for Protein Sciences, Tsinghua University, for the assistance of using the confocal microscopy. This work was supported by the National Natural Science Foundation of China (22477066 and 92478128 to W.Q.; 22407141 to Y.C.; 2240070422 to Y.W.; 32571031 to C.Z.), National Key Research and Development Program of China (No. 2024YFA1308000, W.Q.), Natural Science Foundation of Beijing (JQ25018, W.Q.), Chinese Academy of Medical Sciences (CAMS) Innovation Fund for Medical Sciences (2024-I2M-3-010, Y.C.), Youth Talent Cultivation Fund of Tsinghua University (W.Q.), “Dushi Plan” from Tsinghua University (W.Q.; C.Z.), Beijing Frontier Research Center for Biological Structure, the Fundamental Research Funds from Beijing National Laboratory for Molecular Sciences (BNLMS202301, W.Q.) and the Shenzhen Medical Research Fund (B2401004, W.Q.), China Postdoctoral Science Foundation (No. 2024M761619, Y.W.). W.Q. is supported by Bayer Investigator Award. The proteomic data of human brain from the Banner cohort are based on data obtained from the AMP-AD Knowledge Portal (https://adknowledgeportal.synapse.org/), generated in part from samples collected through the Sun Health Research Institute Brain and Body Donation Program of Sun City, Arizona.

## AUTHOR CONTRIBUTIONS

Y.Z.C., Z.L., Y.W., C. Z., Y.C. and W.Q. designed the research. Y.Z.C. performed all experiments and analyzed all the data except where noted. Y.Z.C performed the proteomic experiments with help from Y.W and W.L. Z.L. performed the functional research for SGK3 and the proteomic experiments of cell surface RTPP. Y.W. performed the proteomic experiments of PL-RTPP and AD samples, and performed RIP-seq. H.L. performed the bioinformatics data analysis for RIP-seq. Y.Z.C, Z.L, C. Z., Y.C. and W.Q. wrote the paper with inputs from all authors.

## SUPPLEMENTAL INFORMATION

**Table S1 Analysis of RRPs in HEK293T identified by RTPP, related to Figure 1 and 2**.

**Table S2 RIP-seq analysis of the RNA clients of HMGCL, PHKG2, and SGK3,related to Figure 2I, 2J, 3B and S3B.**

**Table S3 Analysis of RRPs across all organs identified by RTPP, related to Figure 4**.

**Table S4 Analysis of RRPs in normal and AD hippocampus or other brain region, related to Figure 5**.

**Table S5 Analysis of RRPs in HEK293T cells by PL-RTPP with TurboID-NES and TurboID-NLS, related to Figure 6**.

**Table S6 Development of cell surface RTPP to identify proteins regulated by cell surface RNA, related to Figure 7**.

---

**Figure S1.**
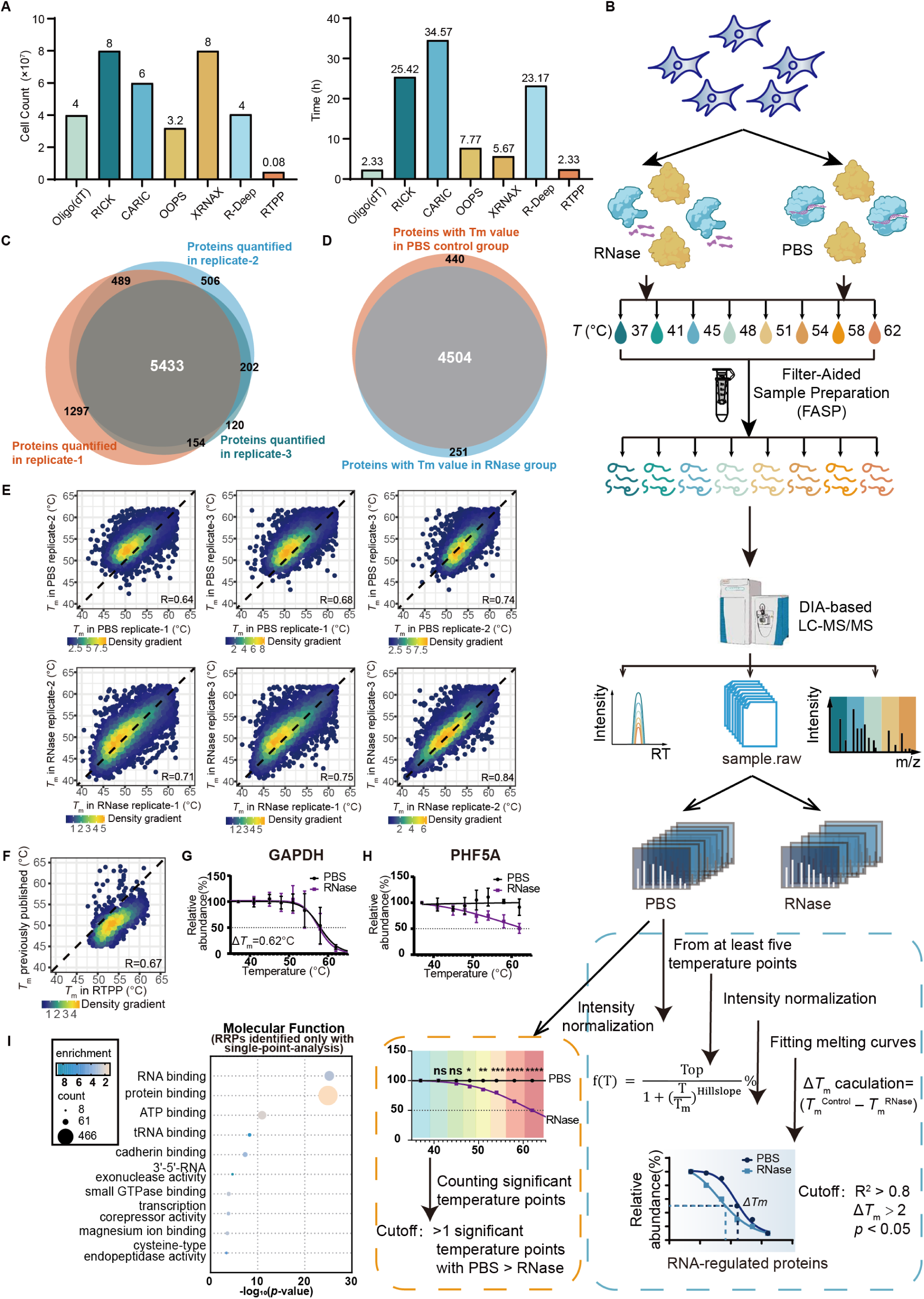
Proteomic profiling of RNA-regulated proteins via data-independent acquisition (DIA)-based RTPP, related to Figure 1. (A) Comparison of starting materials and processing time required for RTPP and other RBP profiling methods. (B) Schematic of the DIA-based streamlined workflow and data analysis process for RTPP proteomics. Cell lysates were treated separately with PBS and RNase, then divided into eight aliquots and heated at the indicated temperatures. After centrifugation, the supernatants were processed into peptides using the FASP workflow. Peptides were then analyzed by LC-MS/MS in DIA mode and quantified using DIA-NN. The raw DIA-based RTPP proteomic data were processed to identify and quantify proteins, followed by a series of data filtering steps, including temperature point verification, intensity normalization, melting curve fitting, and Δ*T*_m_ calculation. Another method of analysis by comparing the significance point by point was also demonstrated. (C) Venn diagram showing the number of quantified proteins across three replicates. (D) Venn diagram showing the number of proteins quantified in both RNase-treated and PBS-treated conditions. (E) Correlation of *T*_m_ values across biological replicates in RNase-treated and PBS-treated conditions. (F) Comparison of *T*_m_ values under normal conditions, quantified by RTPP, against a previous study in HEK293T cells. (G) Thermal melting curve of non-RBP GAPDH. Δ*T*_m_ value was calculated using python. (H) Thermal melting curve of a representative super stable protein with all seven temperature points significantly higher in PBS group than in RNase group. In (G) and (H), error bars represent the mean ± s.d. (I) GO-Molecular Function analysis of RRPs only identified with single temperature point analysis.

**Figure S2.**
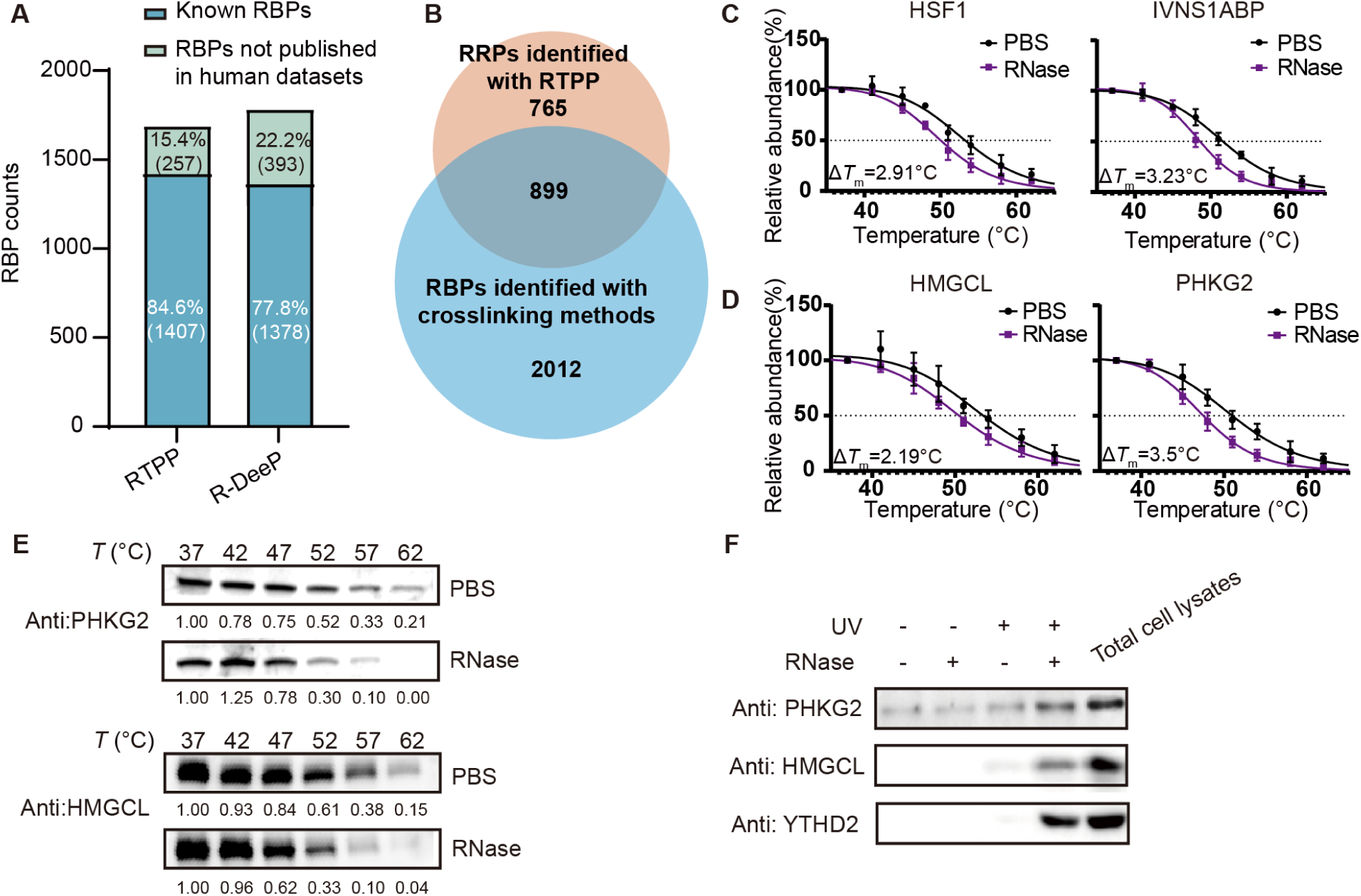
Additional analysis and validation of RNA-regulated proteins identified by RTPP, related to Figure 2. (A) Number and percentage of known RBPs identified by RTPP and R-DeeP. (B) Overlap between RTPP-identified RRPs and RBPs identified by crosslinking-based RBP profiling methods in HEK293T cells. (C) Thermal melting curves of HSF1 and IVNS1ABP determined by DIA-based RTPP. (D) Thermal melting curves of HMGCL and PHKG2 determined by DIA-based RTPP. In (C) and (D), error bars represent the mean ± s.d. (E) Western blot validation of HMGCL and PHKG2 by RTPP (F) Western blot validation of HMGCL and PHKG2 by OOPS

**Figure S3.**
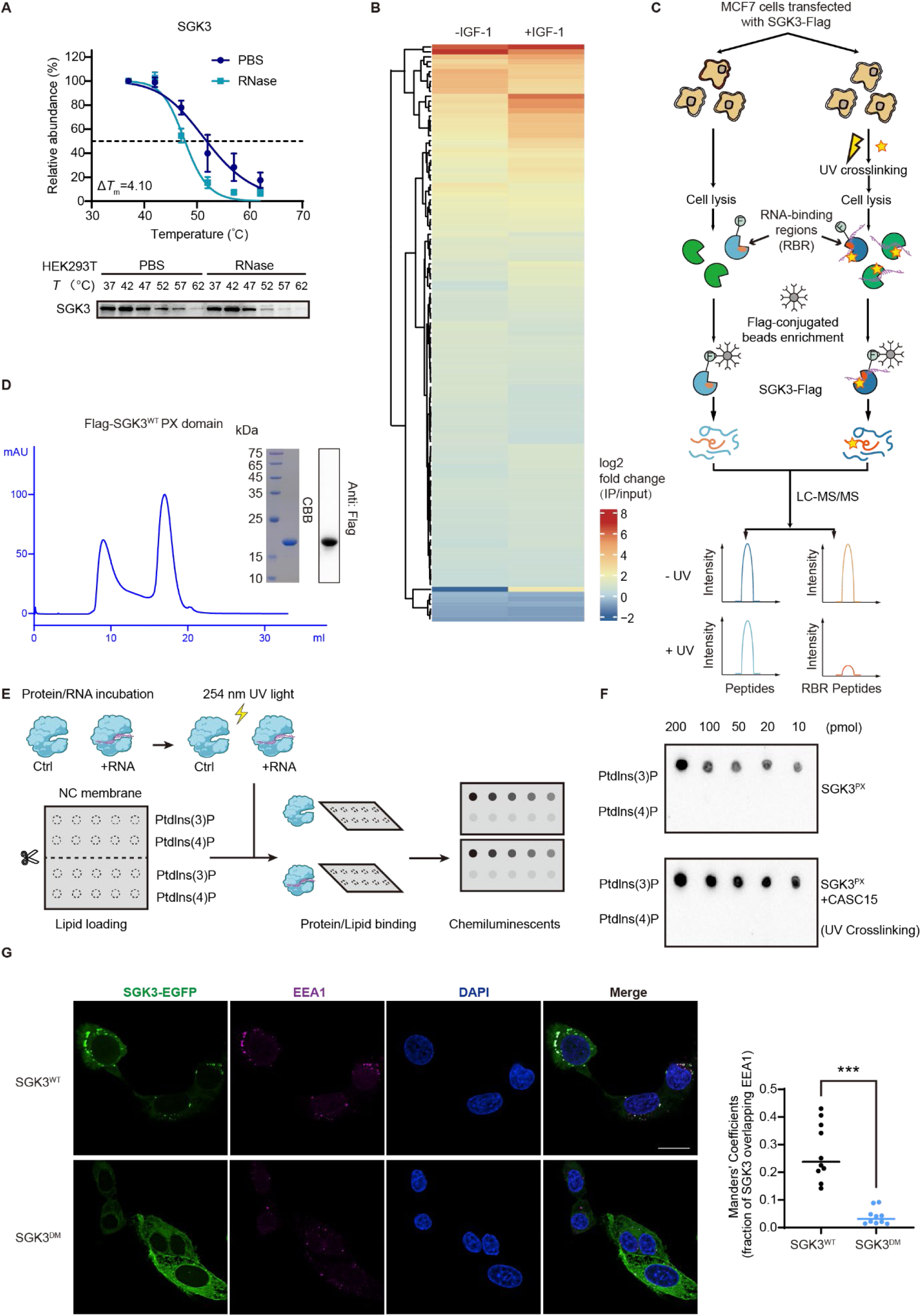
CASC15 promotes SGK3 kinase activity by facilitating its PtdIns(3)P binding and endosomal recruitment, related to Figure 3. (A) Thermal melting curves of SGK3 determined by Western blot. Error bars represent the mean ± s.d. (B) Heatmap showing SGK3-associated RNAs under basal and IGF-1 treatment conditions. (C) Workflow of RBR-ID for finding interaction domain of SGK3. (D) Purity of Flag-SGK3 PX domain. The UV absorption curve was shown with flow volume. Coomassie brilliant blue (CBB) staining and anti-Flag blotting were shown. (E) Workflow of lipid overlay assay for investigating the binding affinity of SGK3 PX domain towards Ptdlns(3)P and Ptdlns(4)P. (F) Dot blot detection of SGK3 lipid binding. 10 nM SGK3 PX domain were incubated with 10 nM CASC15 for 1 hour, followed by UV crosslinking and incubation with nitrocellulose membrane pre-coated with various amounts of Ptdlns(3)P or Ptdlns(4)P. (G) Immunofluorescence of GFP-tagged SGK3^WT^ and SGK3^DM^, co-stained with the endosome marker EEA1 in U2OS cells. The fraction of EGFP overlapping with endosomes were quantified from 10 cells for each experimental condition.\*\*\**p* < 0.001 by ordinary one-way ANOVA. Scale bars, 20 μm.

**Figure S4.**
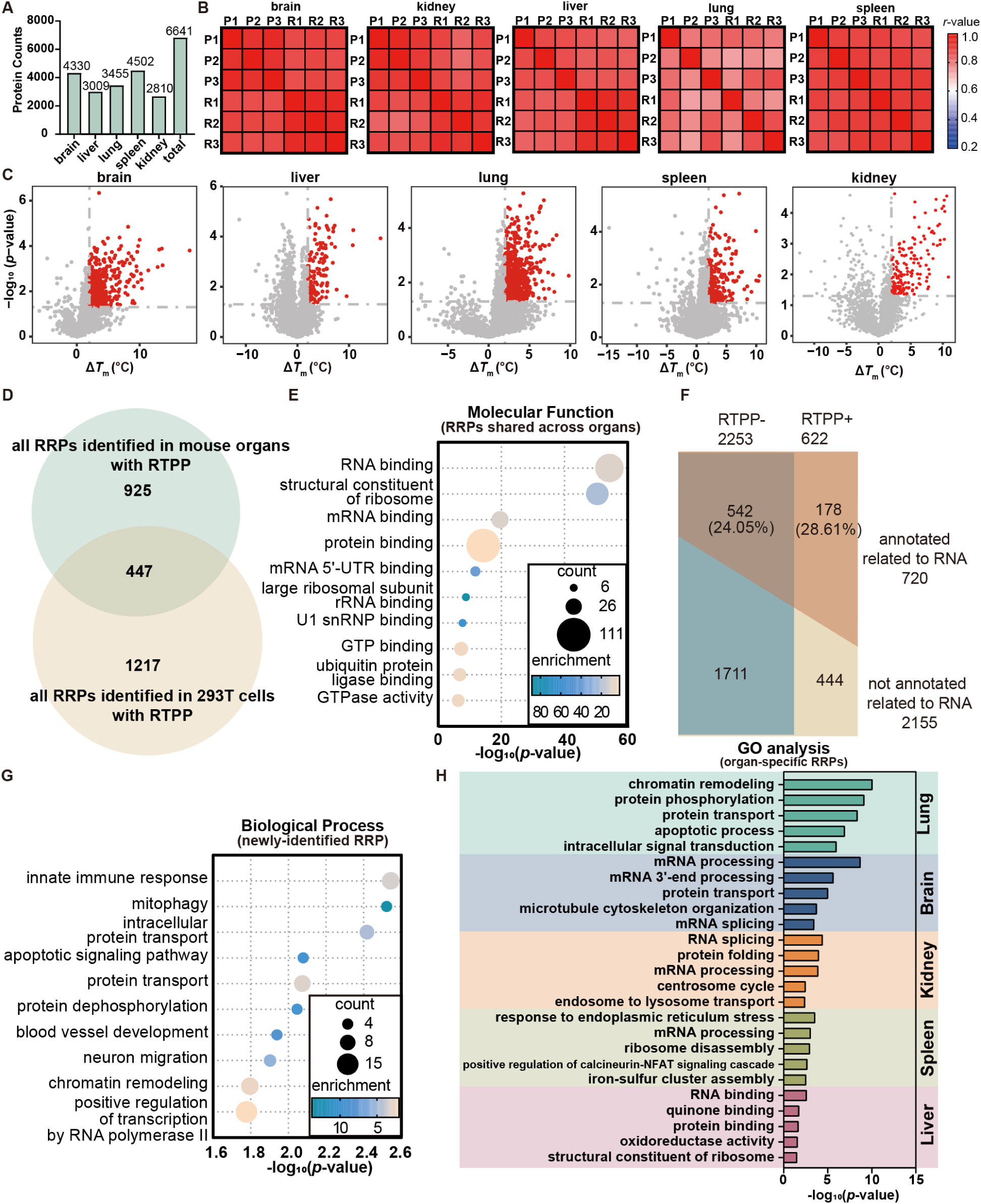
Identification of RNA-regulated proteins across mouse organs, related to Figure 4. (A) Number of proteins quantified in five organs. (B) Correlation matrix of protein intensities among six groups in each organ. The color of each square represents the r value between the 2 groups. (C) Volcano plot of proteins quantified with Δ*T*_m_ by RTPP from five mouse organs. Proteins with Δ*T*_m_ > 2°C and *p*-value < 0.05 were annotated as RRPs and labeled in red. *P*-value was calculated with a two-sided student’s t-test. (D) Venn diagram showing the overlap between RRPs identified in HEK293T cells and all RRPs identified in mouse organs. (E) GO-Molecular Function analysis of RRPs identified in more than one organ. (F) Division of RBPs identified in crosslinking-based methods in mouse by whether identified by RTPP in mouse organs. The functionality of RBPs identified is defined by annotation in GO. In each case, percentage of functional RBPs in RTPP-identified group and RTPP-unidentified group were separately calculated. (G) GO-Biological Process analysis of RRPs in mouse organs previously not identified as RBP in mouse. (H) GO analysis of organ-specific RRPs respectively. GO-Biological Process analysis for lung, brain, kidney, spleen, and GO-Molecular Function analysis for liver were shown.

**Figure S5.**
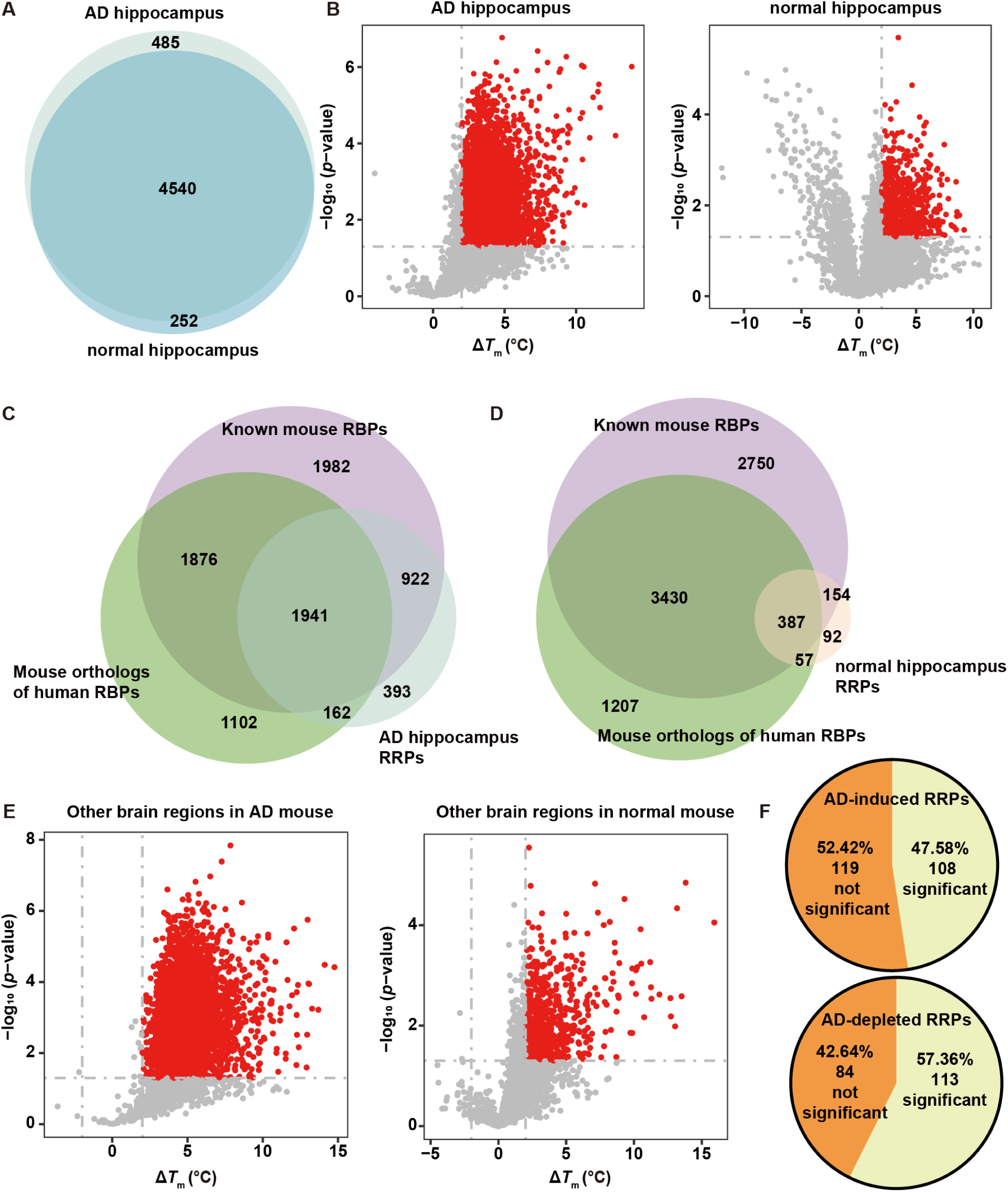
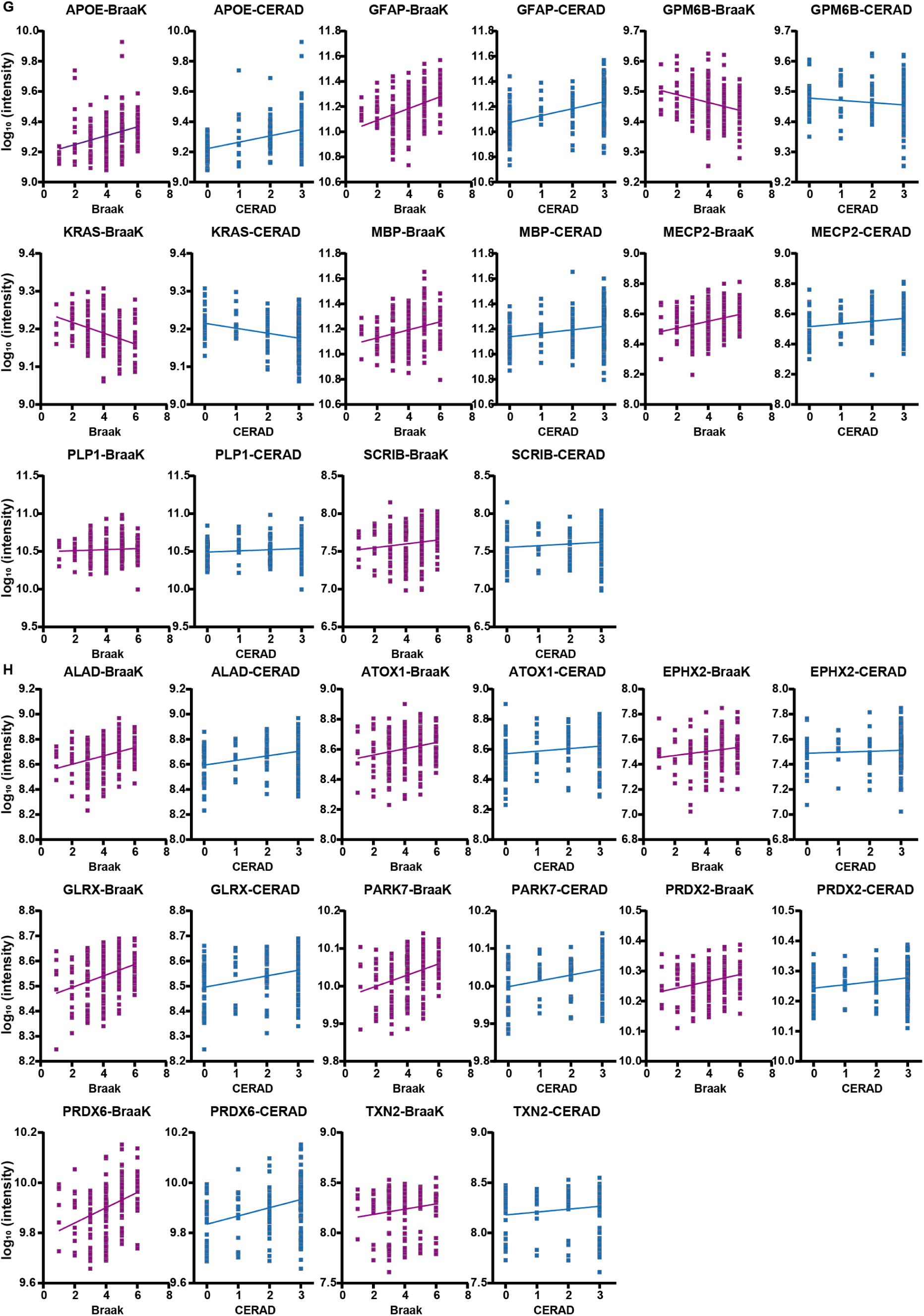
Identification of Alzheimer’s disease-relevant RNA-regulated proteins in mouse hippocampus, related to Figure 5. (A) Total number of proteins quantified in both normal and AD hippocampi. (B) Volcano plot of proteins quantified with Δ*T*_m_ by RTPP in normal and AD hippocampi. Proteins with Δ*T*_m_ > 2°C and *p*-value < 0.05 were annotated as RRPs and labeled in red. (C) Overlap between RRPs identified in the normal hippocampus, known mouse RBPs, and mouse orthologs of human RBPs. (D) Overlap between RRPs identified in the AD hippocampus, known mouse RBPs, and mouse orthologs of human RBPs. (E) Volcano plot of proteins quantified with Δ*T*_m_ by RTPP in other brain regions of normal and AD mice. Proteins with Δ*T*_m_ > 2°C and *p*-value < 0.05 were annotated as RRPs and labeled in red. (F) Proportions of AD-induced and AD-depleted RRPs with significant differences in protein intensities between normal people and AD patients. (G) Protein intensities of representative AD-induced RRPs in patients with different CERAD and Braak scores. (H) Protein intensities of representative AD-depleted RRPs in patients with different CERAD and Braak scores.

**Figure S6.**
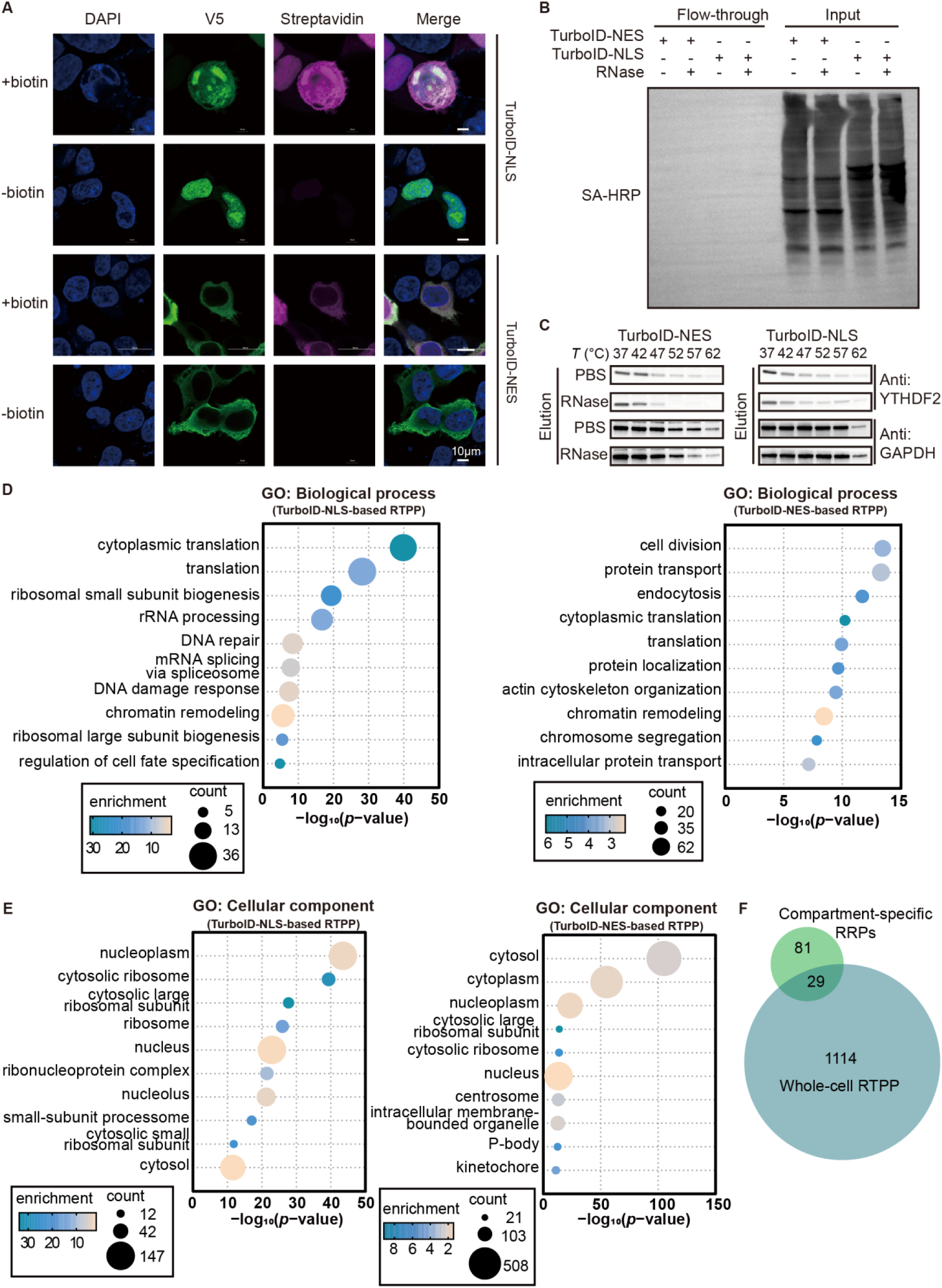
Validation of PL-RTPP and additional analysis of subcellular RNA-regulated proteins identified by PL-RTPP, related to Figure 6. (A) Fluorescent Imaging of TurboID labeling. HEK293T cells transfected with TurboID-NLS and TurboID-NES were treated with 50 μM biotin for 30 minutes. Streptavidin staining indicates biotin labeling, and anti-V5 staining indicates TurboID expression. Scale bars, 10 μm. (B) Streptavidin blotting of total cell lysates (input) and the supernatant after streptavidin enrichment (flow-through). (C) Western blotting of YTHDF2 and GAPDH in eluates of streptavidin enrichment. (D) GO-Biological Process analysis of RRPs identified by TurboID-NLS-based RTPP (left) and TurboID-NES-based RTPP (right). (E) GO-Cellular Components analysis of RRPs identified by TurboID-NLS-based RTPP (left) and TurboID-NES-based RTPP (right). (F) Overlap of compartment-specific RRPs and RRPs identified in whole-cell RTPP.

**Figure S7.**
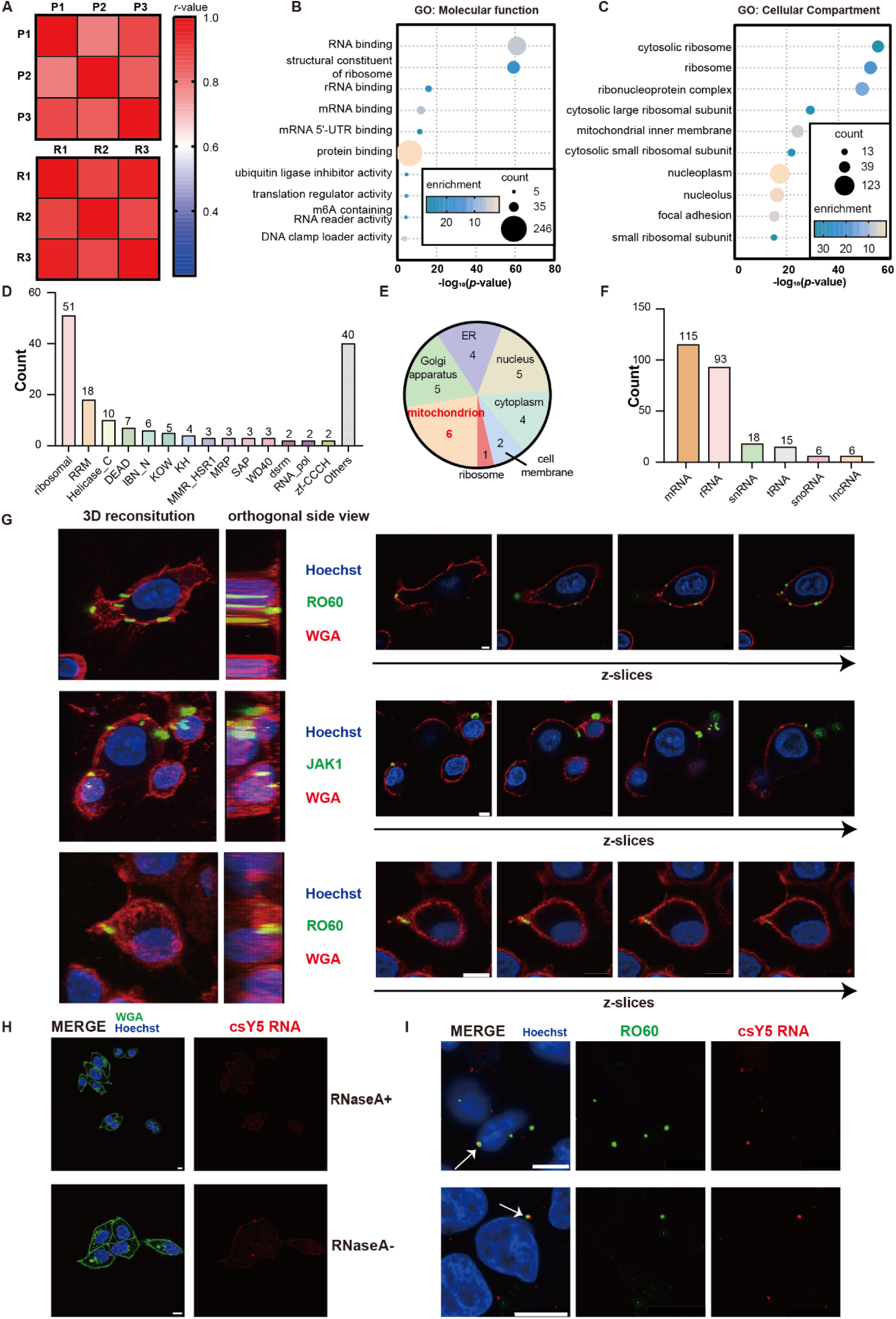
Development of cell surface RTPP reveals nanoscale clusters of RNA-protein complexes on living cells, related to Figure 7. (A) Correlation matrix of three PBS-treated replicates and their corresponding RNase-treated replicates in csRTPP. The color of each square represents the r value between the two groups. (B) GO-Molecular Function analysis of csRRPs. (C) GO-Cellular Components analysis of csRRPs. (D) Subclassification of csRRPs identified by csRTPP with their RNA binding domain profiles. (E) Cellular compartment distribution of the 27 orphan csRRPs identified. (F) Subclassification of csRRPs identified by csRTPP with their associated RNA types. (G) Confocal microscopy z-stack images of candidate csRRPs, co-stained with WGA and Hoechst33342 on live cells. Scale bar, 10 μm. (Left) 3D reconstitution with a slight rotation angle. (Middle) Side view of the 3D reconstitution. (Right) Four z-slices showing distribution of candidate csRRPs on cell surface. (H) Confocal microscopy images showing the effect of RNase A treatment in cleaving cell surface Y5 RNA. Scale bar, 10 μm. (I) Confocal microscopy images of RO60, co-stained with Y5 RNA FISH on cell surface, along with Hoechst33342 on live cells. Scale bar, 10 μm.

## STAR★METHODS

### KEY RESOURCES TABLE

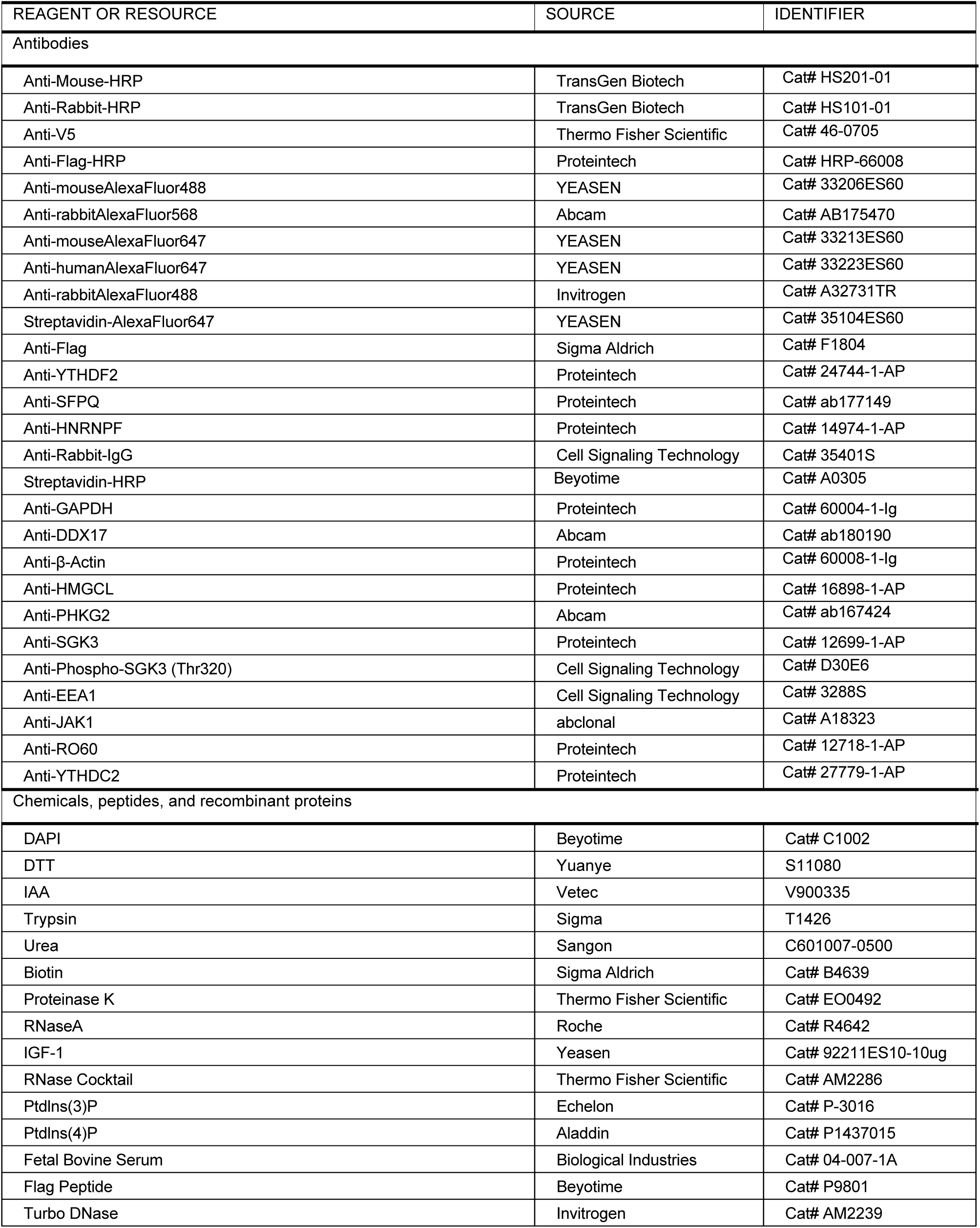
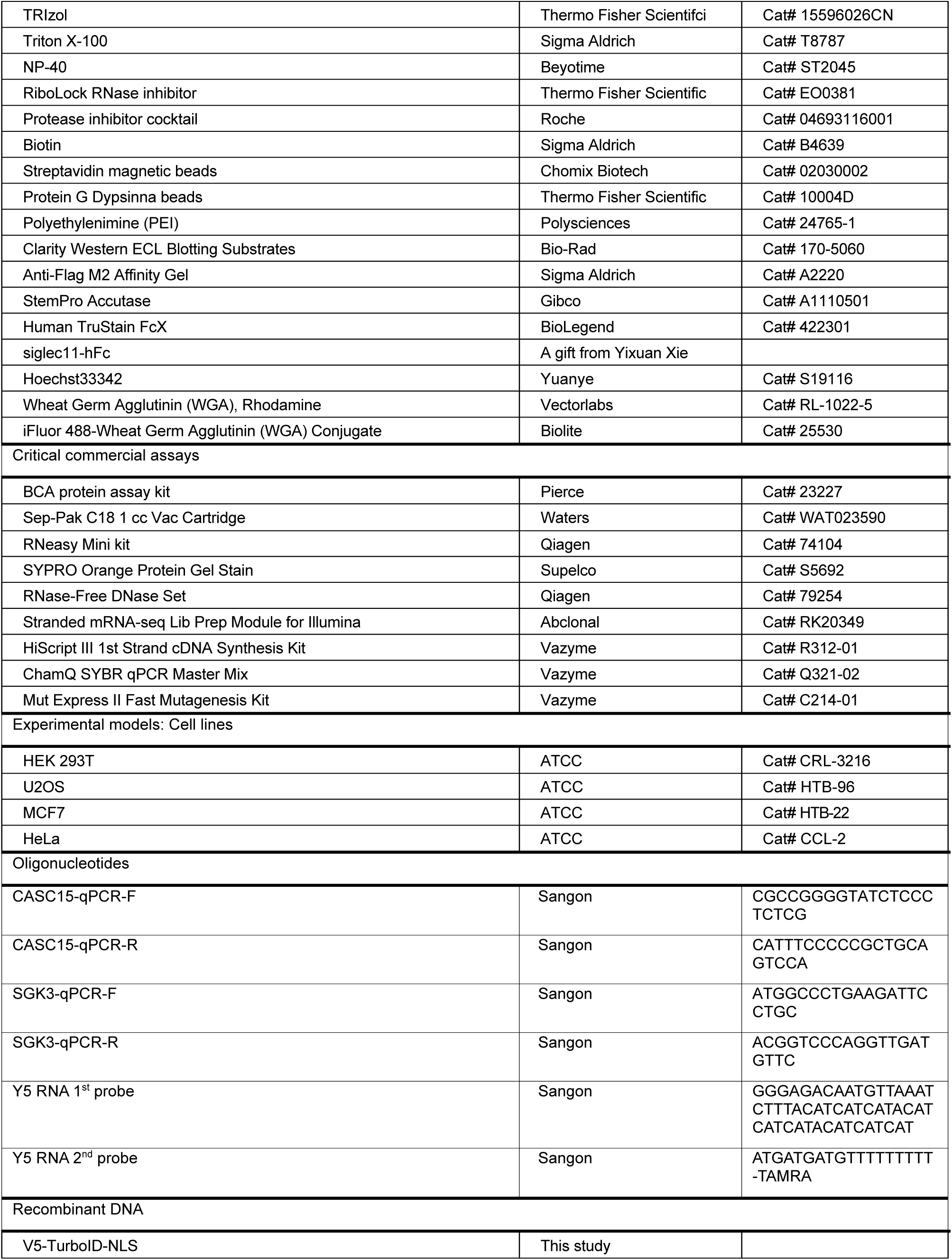
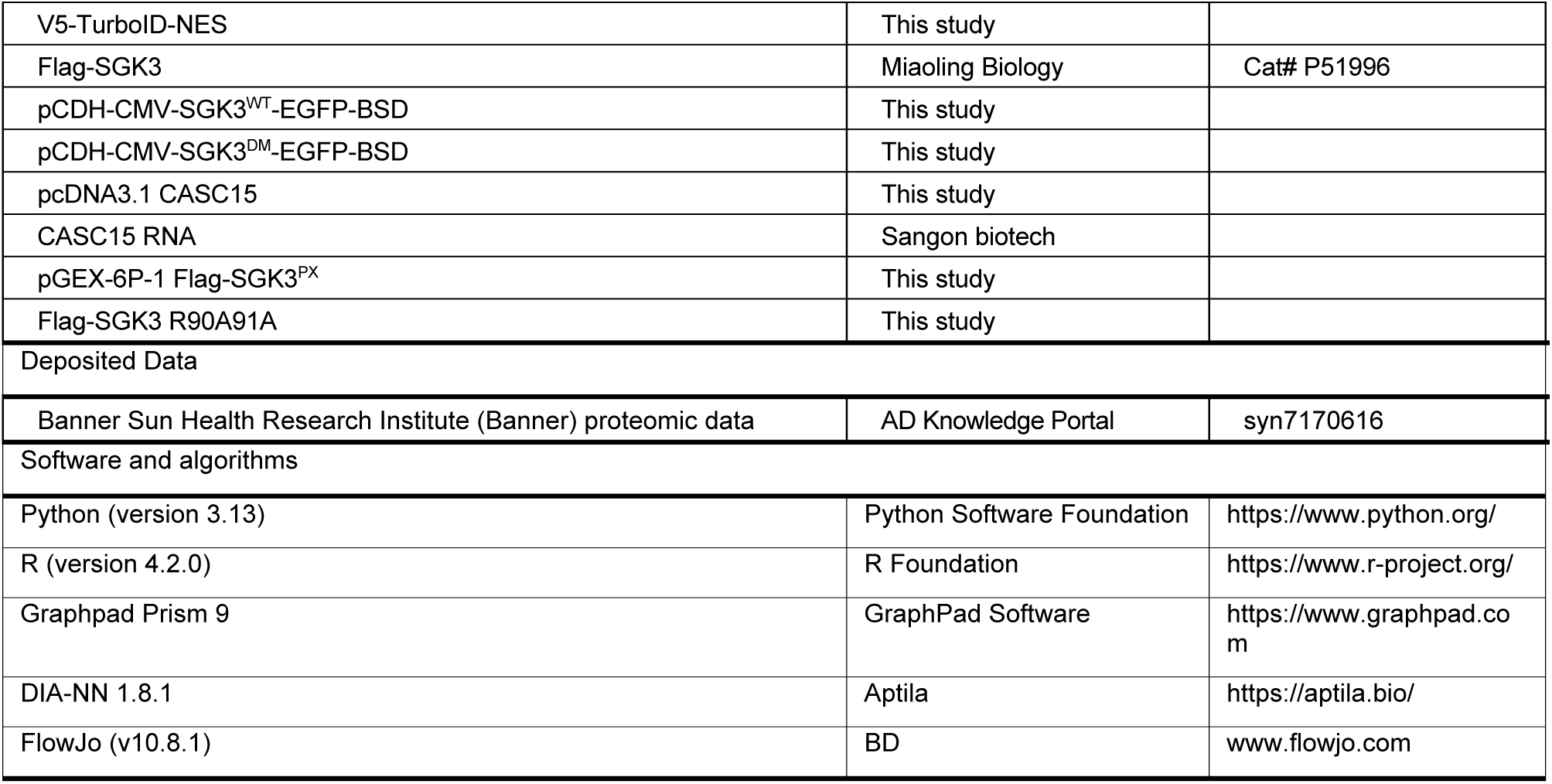

### EXPERIMENTAL MODEL AND STUDY PARTICIPANT DETAILS

#### Cell culture and animals

HEK293T cells, HeLa cells, U2OS cells, and MCF7 cells from the ATCC (passages <25) were cultured in Dulbecco’s modified Eagle’s medium (Gibco by Life Technologies) supplemented with 10% FBS (Biological Industries), 100 U/ml penicillin, and 100 mg/mL streptomycin. Complete medium is supplemented with 10% (vol/vol) fetal bovine serum and 1% (vol/vol) penicillin-streptomycin. All cell lines were cultured in a humidified atmosphere at 37 °C with 5% CO2.

For fluorescence microscopy imaging experiments, cells were grown on 35mm dishes with glass bottom(Cellvis). For Western blotting experiments, RBR-ID and OOPS assays, cells were grown on 10-cm dishes (NEST). For RIP-seq assays, cells were grown on 6-well plates. For proteomic analysis, cells were grown on 15-cm dishes.

5xFAD transgenic mice were maintained by crossing WT mice (C57BL/6J background) with 5xFAD mice (B6SJL background, #034840-JAX, The Jackson Laboratory, Bar Harbor, United States). Female mice were used, and non-transgenic WT littermate mice served as control. Animals were housed under standard conditions of (23 ± 1 °C) and 12-h light/dark cycle with free access to water and food. Animal care and handling were performed according to terms approved by the Institutional Animal Care and Use Committee of Beijing Institute of Technology (Beijing, China).

### METHOD DETAILS

#### RTPP in cells and tissues

For RTPP analysis in HEK293T cells in Figure 1 and Figure S1, HEK293T cells were cultured in 15-cm-diameter dishes until reached 90% of confluence. The cells were washed twice with PBS and harvested using a scraper. The cells then were resuspended in 1 mL of PBS containing EDTA-free protease inhibitor cocktails.

For RTPP analysis across various tissue organs in Figure 3, 150 mg of lung, liver, spleen, kidney and brain were dissected for a single proteomic sample. Organs were washed twice with PBS and cut into small pieces (1-2 mm). The tissues were homogenized 50 times by using a tissue homogenizer containing 1 mL of cold PBS (containing EDTA-free protease cocktail). The resulting homogenized tissue suspension was transferred into a 1.5 mL centrifuge tube.

For RTPP analysis of the normal or AD mouse hippocampus and other brain regions in Figure 4, three female 5×FAD transgenic mice were maintained by crossing WT mice (C57BL/6J background) with 5×FAD mice (B6SJL background, #034840-JAX, The Jackson Laboratory, Bar Harbor, United States). Three non-transgenic WT littermate mice served as control. A total of 150 mg of hippocampus and 100 mg of other brain regions were dissected for a single proteomic sample. The tissues were homogenized 50 times by using a tissue homogenizer containing 1 mL of cold PBS (containing EDTA-free protease cocktail). The resulting homogenized tissue suspension was transferred into a 1.5 mL centrifuge tube.

All cell and tissue samples were lysed by snap-freezing in liquid nitrogen. The tube was placed in a thermal mixer at 25°C until most of the contents had thawed, and then transferred to ice until the entire content was fully thawed. The freeze–thaw cycle was repeated five times before the samples were centrifuged for 20 minutes at 4°C and 20,000 × *g*. The supernatant was collected and protein concentration was determined using the Pierce BCA protein assay kit. 40 μL of RNase cocktail (containing 20 U of RNaseA and 800 U of RNase T1) or PBS was added to cell lysates containing 3 mg of proteins with protein concentration of 3mg/mL and incubated for 1 hour at 4 °C. Cell lysates were divided into eight aliquots (100 μL for each aliquot) and transferred into 0.2-mL PCR tubes. Each sample was heated in parallel for 3 minutes to the respective temperature (37 °C, 41 °C, 45 °C, 48 °C, 51 °C, 54 °C, 58 °C and 62 °C). Subsequently, the samples were centrifuged at 20,000 × *g* for 20 minutes at 4 °C and the supernatants were collected.

#### RTPP for western blotting

For Western blotting-based RTPP, samples were snap-frozen in liquid nitrogen. The samples were lysed by snap-freezing in liquid nitrogen. The tube was then placed in a thermal mixer at 25°C until most of the contents had thawed, after which it was transferred to ice until fully thawed. The freeze–thaw cycle was repeated five times before the samples were subjected to centrifugation for 20 minutes at 4°C and 20,000 × *g*. The supernatant was collected, and protein concentration of the supernatant was determined using the Pierce BCA protein assay kit. 40 μL of RNase cocktail (containing 20 U of RNaseA and 800 U of RNase T1) or PBS was added to cell lysates containing 2 mg of proteins with protein concentration of 2mg/mL and incubated for 1 hour at 4 °C. Cell lysates were divided into six aliquots (50 μL, 2 mg/mL for each aliquot) and transferred into 0.2-mL PCR tubes. Each sample was heated in parallel for 3 minutes to the respective temperature (37 °C, 42 °C, 47 °C, 52 °C, 57 °C and 62 °C). Subsequently, the samples were centrifuged at 20,000 × *g* for 20 minutes at 4 °C and the supernatants were collected. 40 μL of supernatants were boiled with 10 μL of 5 × protein loading buffer for subsequent Western blotting.

#### DIA-MS sample preparation after RTPP

For proteomic analysis, Filter-Aided Sample Preparation (FASP) was performed as previously described^35^. 50 μL of the collected supernatant after the RTPP procedure was diluted with 150 μL of PBS containing 8 M urea. 10 mM dithiothreitol (DTT) was added, and the mixture was incubated at 35°C for 30 minutes. Then, 20 mM iodoacetamide (IAA) was added, and the sample was incubated for another 30 minutes at 37°C in the dark. A 10K filter (Millipore) was equilibrated with 300 μL of 10 mM ammonium bicarbonate solution (ABC, pH 10). Each sample was transferred to the 10K filter and centrifuged at 15,000 × g for 5 minutes at 16°C. An additional 200 μL of ABC solution was added to the 10K filter, and the centrifugation and addition cycle was repeated five times. 2 μg of trypsin was added to the proteome sample, and the proteome samples in the 10K filter were digested for 16 hours at 37°C. After digestion, the samples were centrifuged at 20,000 × g for 10 minutes. The remaining material on the filter was washed twice by adding 200 μL of ABC solution and centrifuging at 20,000 × g for 10 minutes. The combined extracts were dried in a vacuum concentrator. The resulting peptide mixtures were reconstituted in 0.1% (v/v) formic acid (FA) in water prior to mass spectrometry analysis.

#### PL-RTPP

For spatiotemporally resolved RTPP in Figure 6 and Figure S6, HEK293T cells were plated in 15-cm dishes and transiently transfected with 10 μg of TurboID-NES or TurboID-NLS for 24 hours. TurboID labeling was initiated by adding a final concentration of 50 μM biotin for 30 minutes. After labeling, cells were washed three times by PBS. The cell lysates were then subjected to the cell-based RTPP procedure. To enrich biotinylated proteins, 200 μL of streptavidin-coated magnetic beads (Chomix Biotech) were washed twice with RIPA lysis buffer (50 mM Tris pH 8, 150 mM NaCl, 0.1% SDS, 0.5% sodium deoxycholate, 1% Triton X-100, 1 × protease inhibitor cocktail, and 1mM PMSF), then incubated with 1 mL of collected supernatants in RIPA buffer with rotation at 4 °C overnight. The beads were subsequently washed twice with 1 mL of RIPA lysis buffer, once with 1 mL of 1 M KCl, once with 1 mL of 0.1 M Na_2_CO_3_, once with 1 mL of 2 M urea in 10 mM Tris-HCl (pH 8.0), and twice with 1 mL of RIPA lysis buffer. For Western blot analysis, the enriched proteins were eluted by boiling the beads in 100 μL of 1× protein loading buffer supplemented with 20 mM DTT and 2 mM biotin. For proteomic analysis, the beads were washed twice with 1 mL of PBS, then resuspended in 200 μL of 50 mM Tris-HCL buffer (pH 7.5), transferred into new 1.5 mL Eppendorf tubes.

For on-bead trypsin digestion, streptavidin magnetic beads post enrichment were washed twice with 200 μL of 50 mM Tris-HCl buffer (pH 7.5). The Tris-HCl buffer was removed, and beads were incubated in 0.4 μg trypsin in 80 μL of 2 M urea/50 mM Tris buffer with 1 mM DTT, for 1 hour at room temperature while shaking at 1150 rpm. Following pre-digestion, 80 μL of each supernatant was transferred into new tubes. Beads were then washed twice with 60 μL of 2 M urea in 50 mM Tris (pH 7.5) buffer, and these washes were combined with on-bead digest supernatant. The eluates were spun down at 5,000 × g for 30 seconds and the supernatant was transferred to a new tube. Samples were reduced with 4 mM DTT for 30 minutes at 35 °C with 800 rpm of shaking. Following reduction, samples were alkylated with 10 mM iodoacetamide for 30 minutes in the dark at 35 °C with 800 rpm of shaking. An additional 0.5 μg of trypsin was added and samples were digested overnight at room temperature while 800 rpm of shaking. Following overnight digestion, samples were desalted on C18 StageTips (Waters). The C18 Tip was activated once with 1 mL of methanol, once with 1 mL of acetonitrile and once with 1 mL of 50% (v/v) acetonitrile and 0.1% (v/v) formic acid (FA), and then was equilibrated three times with 1 mL of 0.1 % (v/v) FA. The digested samples were transferred to C18 Tip and loaded three times. The samples with C18 Tip were washed 10 times with 1 mL of 0.1% (v/v) FA. The peptides were eluted three times with 300 μL of 50% (v/v) acetonitrile and 0.1% (v/v) FA to a new 1.5 mL Eppendorf tube. Eluted peptides were dried in a vacuum concentrator to completion and stored at −80°C.

#### Cell surface RTPP

For resolving cell surface RTPP in Figure7, HEK293T cells were plated in 15 cm dishes until reached 90% of confluence. The intact cells were washed twice with PBS and treated with PBS or 0.1 mg/mL RNase A (Roche) at 37 °C for 10 min. Then the cells were washed once with PBS and harvested using a scraper. The cells then were resuspended in 500 μL of PBS containing 0.4 % NP-40 and EDTA-free protease inhibitor cocktails. The samples were lysed by snap-freezing in liquid nitrogen. The tube was then placed in a thermal mixer at 25°C until most of the contents had thawed, after which it was transferred to ice until fully thawed. The freeze-thaw cycle was repeated five times before the samples were subjected to centrifugation for 20 minutes at 4°C and 20,000 × g. The supernatant was collected, and protein concentration of the supernatant was determined using the Pierce BCA protein assay kit. Cell lysates were divided into six or eight aliquots (50 μL, 2 mg/mL for each aliquot) and transferred into 0.2-mL PCR tubes. Each sample was heated in parallel for 3 minutes to the respective temperature. Subsequently, the samples were centrifuged at 20,000 × g for 20 minutes at 4 °C and the supernatants were collected. 40 μL of supernatants were boiled with 10 μL of 5 × protein loading buffer for subsequent Western blotting. For proteomic analysis, Filter-Aided Sample Preparation (FASP) was performed as described in DIA-based RTPP.

#### DIA-based LC-MS/MS analysis

The loading and separation of peptides were achieved using a reversed-phase C18 column (trap column: 5 cm × 100 μm, 3 μm particle size, analytical column: 20 cm × 100 μm, 1.9 μm particle size). The mobile phases (A: water with 0.1% formic acid, B: 94% acetonitrile with 0.1% formic acid) were driven and controlled by a Vanquish™ Neo UHPLC system (Thermo Fisher Scientific) at flow rate of 300 nL/minutes. Peptides were chromatographically separated using a 166-minutes gradient from 5% to 60% solvent B. The LC gradient for protein samples began with an increased from 4% to 5% for the first 4 minutes of the analysis, followed by an increase from 5 % to 20% B from 4 to 109 minutes, an increase from 20% to 35% B from 109 to 150 minutes, an increase from 35% to 99% B from 150 to 159 minutes, and holding at 99 % for the last 7 minutes to wash the column.

For the samples analyzed by Q Exactive-plus series Orbitrap mass spectrometers (Thermo Fisher Scientific), the precursors were ionized using a HESI Ⅱhigh-energy surface ionization source (Thermo Fisher Scientific) held at +2.2 kV relative to ground, and the inlet capillary temperature was maintained at 320 °C in positive-ion mode. The total measurement time for each sample was 159 minutes. Peptide groups were equally distributed across 45 windows according to their mass-to-charge ratio. The number and width of the isolation windows were manually calculated. MS data were acquired using the data independent acquisition (DIA) mode, with each DIA cycle consisting of one full scan and the calculated number of DIA scans covered a mass range of 350-1800 Th, with an overlap of 1 Th. For the full MS analysis, resolution was set to 70,000 and full MS AGC target was 3E6 with an IT of 20 ms. For each DIA window, resolution was set to 17,500, the AGC target value for fragment spectra was set to 1E6, with an auto IT. The normalized CE was set at 28%. The default charge state was set to 3, and the fixed first mass was set to 200 Th. Other parameters were set to their default values.

#### DIA-NN analysis

For peptide and protein identification and quantification, the raw data were processed using DIA-NN 1.8.12 in an advanced, library-free module. The main search settings for in silico library generation were set as follows: trypsin/P with a maximum 3 missed cleavages, protein N-terminal M excision enabled, carbamidomethyl on cysteine as a fixed modification, oxidation on methionine as a variable modification, peptide lengths ranging from 7 to 30, precursor charge states from +1 to +4, precursor m/z range from 350 to 1800, fragment m/z range from 350 to 1800. Full MS and MS/MS spectra were searched against a concatenated forward and reverse version of the *Homo sapiens* and *Mus musculus* database, downloaded from UniProt (https://www.uniprot.org/).

Additional search parameters were set as follows: the quantification strategy was set to ‘robust LC (high precision)’ mode, cross-run normalization was RT-dependent, MS1 and MS2 mass accuracies were set to 0, allowing DIA-NN to automatically determine mass tolerances, the scan window was set to 6 corresponding to the approximate average number of data points per peak, isotopologues and MBR were enabled, the neural network classifier was set to single-pass mode.

#### Normalization of DIA-based quantification data

The data analysis of RTPP was performed as described previously^25^. For DIA-based quantification, the intensities for eight temperature points were collected for each protein. The relative intensity values were then calculated by using the maximum intensity corresponding to the two lowest temperatures (37 °C and 42 °C) as the reference, scaled on a 0–100% scale. Proteins that had more than one unique quantified peptide were selected for further analysis.

For data preprocessing, three assumptions were made: (1) the whole proteome remained stable at temperatures lower than 37 °C and completely unfolded at temperatures proximal to 100 °C, (2) RNase did not affect the thermal stability of the whole proteome, such that the global melting curves were similar between PBS-treated and RNase-treated proteomes, and (3) the intensity of the reporter ions at any given temperature point never exceeds the ion intensities at the two adjacent temperature points. As a result, two additional data points, 0% at 97 °C and 100% at 27°C, was added to the relative intensities for each protein to fit the melting curve after normalization by the corresponding correction factors, in order to reduce global drifting at each temperature. At least four temperature points with DIA quantification intensities were required for a unique quantified peptide to ensure accurate fitting of the melting curve. If the intensity at any temperature point was missing, the information from the next higher temperature point was used as a substitute for fitting melting curves (see ‘Fitting melting curves’ for details). For each group of three replicate samples either treated with PBS or with RNase, a set of eight correction factors were calculated, with each factor corresponding to a specific temperature point in the RTPP experiment. First, the relative DIA intensities of all the identified proteins at each of the eight temperature points were averaged to fit a global melting curve for the PBS-treated or RNase-treated sample. Second, for the temperature point with the largest fitting error in the global melting curve, its averaged DIA intensity was adjusted by the corresponding correction factor, aiming to minimize the fitting error and refit the global melting curve. Third, the procedure was iterated multiple times until no further improvement could be gained, resulting in the final set of eight correction factors. Finally, these correction factors were applied to each protein to normalize the relative intensities at the corresponding temperature points so that the melting curve could be fitted. A similar strategy was previously implemented^25^.

#### Fitting melting curves

Normalized relative intensity values as a function of temperature were fitted to a sigmoidal curve according to the following equation using the curve fit function from SciPy^84^.

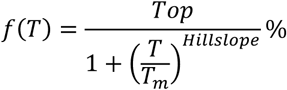

Where HillSlope represents the sharpness of the melting curve. The *T*_m_ of a protein is defined as the temperature at which half of the protein has been denatured. The *T*_m_ value for each protein was calculated using this equation, and only *T*_m_ values derived from melting curves with an *R*^2^ > 0.8 were used in subsequent analyses.

Shifts in *T*_m_ values (Δ*T*_m_) induced by RNase were determined by subtracting the *T*_m_ of the RNase-treated protein from the *T*_m_ of the PBS-treated protein:

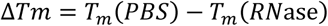

Three biological replicates were performed for RTPP profiling. Considering that some proteins were only partially unfolded (<50% unfolded) at the highest temperature (62 °C) in our profiling, the calculated *T*_m_ values for these proteins might not be accurate. To minimize such inaccuracies, all RRP candidates must meet the following criteria: average*T_m_*(*PBS*) < 62°C and average*T_m_*(*RN*ase) < 62°C.

#### Single temperature point analysis

For single temperature point analysis of RTPP proteomic data from HEK293T cells in Figure S1B, S1H, and S1I, all protein intensities were scaled with its correction factor, and the percentage was calculated to the corresponding protein intensity at 37°C. For a certain protein, all 3 percentages at a temperature point were calculated for average, and a p-value for student’s t-test between PBS group and RNase group at this temperature can be calculated. A protein is identified as an RRP through this method needs to fit the following criteria: (1) More than 1 temperature points have the property that *average(PBS) > average(RNase)*; (2) More than 1 of these points have the property that p-value < 0.05.

#### Analysis of Proteomic Data

After calculating the *T*_m_ for each quantified protein, filtering criteria of Δ*T*_m_ > 2°C and p-value < 0.05 were applied to define RRPs, consistent with the criteria used in previous thermal proteome profiling studies^25,29^. The *p*-value was calculated using a two-sample t-test to assess statistical significance. Gene Ontology (GO) analysis and Kyoto Encyclopedia of Gene and Genomes (KEGG)-pathway analysis in Figure1F, S1I, 2H, 4D, 4E, S4E, S4G, S4H, 5G, 5H, 6H, S6D, S6E, 7D, S7B, and S7C were performed using the DAVID software (https://david.ncifcrf.gov/), which facilitated the identification of enriched biological processes, molecular functions, cellular components, tissue expression and KEGG pathway.

Heatmaps in FigureS3B, 4F, 4J, 5B, and 6E were generated using the complexHeatmap package by R (version 4.2.0). The RNA-binding domains analysis in Figure 2D, 2E, 2F, and S7D was performed using the Pfam database (now hosted by Interpro), The RNA-binding protein types in Figure2B, 2C, 2F, S4F, and S7F were assessed based on the “RNA binding” category from the GO analysis. Protein–protein interaction (PPI) networks in Figure4J, 5I, and 5J were constructed using STRING (Search Tool for the Retrieval of Interacting Genes/Proteins) (https://string-db.org/) to explore the functional interactions between the identified RRPs.

Venn diagrams illustrating overlapping protein sets in FigureS1C, S1D, 2A, S2B, 4C, S4D, 5C, 5D, S5A, S5C, S5D, 6F, 6G, S6F, 7E, and 7G were generated using the Biovenn tool available at http://www.biovenn.nl/index.php, and S1C, S1D, and Figure 4G was generated with R. Box plots and correlation matrices were created using GraphPad Prism software (version 9.5.1). For the GO and KEGG pathway bubble plots, the “multiple variables” module in GraphPad Prism was used. Density scatter plots in Figure S1E, S1F, and volcano plots in Figure 1D, S4C, 5F, S5B. S5E, 6B, 7C were generated with R.

#### Western blots

For all Western blots and silver staining gels, samples were resolved on a 10% SDS-PAGE gel. For all Western blots, after SDS-PAGE, the gels were transferred to a PVDF membrane. The blots were then blocked in 5% (w/v) skim milk in TBS-T (Tris-buffered saline, 0.1% Tween 20) for at least 30 minutes at room temperature. For streptavidin blotting in Figure S6B, the blots were stained with 0.3 mg/mL streptavidin-HRP (1:5000, A0305, Beyotime) in TBS-T for 1 hour at room temperature.

For determining the thermal stability of RBPs in Figure 1B, the blots were stained with anti SFPQ (1:2000 dilution, ab177149, Proteintech), anti-DDX17 (1:2000 dilution, ab180190, Abcam), anti-HNRNPF (1:2000 dilution, 14974-1-AP, Proteintech) and anti-YTHDF2 (1:2000 dilution, 24744-1-AP, Proteintech) in TBS-T with 5% (w/v) BSA overnight at 4 °C. For determining the thermal stability of non-RBPs in Figure 1C, the blots were stained with anti-GAPDH (1:5000 dilution, 60004-1-Ig, Proteintech) and anti-ACTB (1:5000 dilution, 60008-1-Ig, Proteintech), in TBS-T with 5% (w/v) BSA overnight at 4 °C. For the validation of new-identified RRPs in Figure S2F, the blots were stained with anti-PHKG2 (1:1000 dilution, ab167424, Abcam) and anti-HMGCL (1:1000 dilution, 16898-1-AP, Proteintech), in TBS-T with 5% (w/v) BSA overnight at 4 °C. For determining the thermal stability of YTHDF2 and GAPDH in nucleus or cytoplasm in Figure S6C, the blots were stained with anti-YTHDF2 (1:1000 dilution, 24744-1-AP, Proteintech) and anti-GAPDH (1:5000 dilution, 60004-1-Ig, Proteintech), in TBS-T with 5% (w/v) BSA overnight at 4 °C.

For determining the thermal stability of SGK3 in Figure S3A, the blots were stained with anti-SGK3 (1:1000 dilution, 12699-1-AP, Proteintech)

For determining the thermal stability of cell surface RRPs DDX21, HNRNPU and GAPDH in Figure 7B, the blots were stained with anti-DDX21 (1:1000 dilution, 10528-1-AP, Proteintech), anti-HNRNPU (1:1000 dilution, 14599-1-AP, Proteintech), anti-GAPDH (1:5000 dilution, 60004-1-Ig, Proteintech)

For determining the phosphorylation level of SGK3 by immunoprecipitation in Figure 3J, the blots were stained with anti-SGK3 (1:1000 dilution, 12699-1-AP, Proteintech), anti-phospho-SGK3 (Thr320) (1:500 dilution, D30E6, Cell Signaling Technology), anti-GAPDH (1:5000 dilution, 60004-1-Ig, Proteintech) in TBS-T with 5% (w/v) BSA overnight at 4 °C.

After incubating with the primary antibody, the blots were washed with TBS-T for four times (5 minutes for each wash). For the detection of phosphorylation signal, blots were stained with secondary antibodies in TBS-T with 1% skim milk (w/v) for 40 minutes in room temperature. The other blots were stained with secondary antibodies in TBS-T for 40 minutes in room temperature and then washed four times with TBS-T for 5 minutes each time before to development with Clarity Western ECL Blotting Substrates (Bio-Rad) and imaging on the Tanon 5200 Multi System (Tanon). The intensity of Western blot bands was quantified using ImageJ, the data are presented as means ± SD.

#### RIP-seq

For RIP-seq analysis of HMGCL and PHKG2 in Figure 2I and 2J, HEK293T cells were cultured on 6-well plates. Protein G beads (Thermo Fisher Scientific) were conjugated with antibodies before immunoprecipitation. 40 μL of Protein G beads were washed four times with 450 μL of lysis buffer (50 mM Tris-HCl, pH 7.4, 100 mM NaCl, 1% NP-40, 0.1% SDS, 0.5% sodium deoxycholate, 1% EDTA-free protease inhibitor cocktail in DEPC water). The beads were then resuspended in 200 μL of lysis buffer with 2.5 μg of the HMGCL and PHKG2 antibodies with rotation at 4 °C.

For RIP-seq analysis of SGK3 in Figure 3B, MCF7 cells were cultured on 6-well plates and transiently transfected with 2 μg of Flag-SGK3^WT^ or Flag-SGK3^DM^ for 24 hours. For IGF-1-stimulated conditions, cells were treated with 100 ng/ml of IGF-1 for 20 minutes. Protein G beads were conjugated with antibodies before immunoprecipitation. 40 μL of Protein G beads were washed four times with 450μL of lysis buffer (50 mM Tris-HCl, pH 7.4, 100 mM NaCl, 1% NP-40, 0.1% SDS, 0.5% sodium deoxycholate, 1% EDTA-free protease inhibitor cocktail in DEPC H2O). The beads were then resuspended in 200 μL of lysis buffer with 2.5 μg of the Flag (F1804, Sigma Aldrich) and rabbit-IgG(H/L) (35401S, Cell Signaling Technology) antibody with rotation at 4 °C.

Cells were washed twice with PBS, then recoated with 2 mL of ice-cold PBS (containing EDTA-free protease inhibitor cocktail). The plates, with the lids removed, were placed on ice and crosslinked using a UV crosslinker at 400 mJ/cm^2^. Cells were scraped and collected, then centrifuged at 5,000 rpm for 5 minutes at 4 °C. A total of 2×10^6^ cells were lysed in 150 μL of lysis buffer on ice for 5 minutes. The lysate was treated with Turbo DNase (AM2239, Invitrogen, 0.2 U/mL) at 37 °C for 5 minutes with 1,100 rpm shaking, followed by centrifugation at 14,000 × *g* for 10 minutes at 4 °C. 2 μL of supernatant was retained as the “input”, and the remaining lysate was incubated with antibody-conjugated Protein G beads with rotation for 4 hours at 4 °C

The beads were then washed once with 150 μL of High Stringency Buffer (15 mM Tris-HCl pH 7.5, 5 mM EDTA, 1% Triton X-100, 1% Na-deoxycholate, 0.001% SDS, 120 mM NaCl, 25 mM KCl), once with High Salt Buffer (15mMTris-HCl pH 7.5, 5mMEDTA, 1% Triton X-100, 1% Na-deoxycholate, 0.001% SDS, 1M NaCl) and Low Salt Buffer (15mM Tris-HCl pH 7.5, 5 mM EDTA) with rotation for 10 minutes at 4 °C. Both inputs and beads were resuspended in 50 μL of proteinase K digestion buffer (100 mM Tris-HCl pH 7.5, 100 mM NaCl, 0.2 mM EDTA, 0.1% SDS) containing 5 μL of proteinase K (ST532, Beyotime) and 2.5 μL of RNase inhibitor (EO0381, Thermo Fisher Scientific). The mixture was incubated for 2 hours at 50 °C with shaking at 1,100 rpm.

Following digestion, samples were diluted with 150 μL lysis buffer and extracted with 200 μL of RNA extraction reagent (phenol: chloroform: isoamyl alcohol 25:24:1) by fiercely shaking (at least 50 times) in a heavy phase-lock tubes (WM5-2302830, TIANGEN). The samples were centrifuged at 13200 rpm for 5 minutes. An additional 200 μL of chloroform was added, followed by fiercely shaking for at least 50 times. The samples were centrifuged at 13200 rpm for 5 minutes. The supernatant was transferred to a new LowBind centrifuge tube. DNA was digested by incubating the supernatant with 18 μL of Turbo DNase (AM2239, Invitrogen) at 37 °C for 15 minutes with shaking at 1,100 rpm. The RNA was then purified with Phenol-Chloroform again. The purified RNA was mixed with 20 μL of natrium acetate (3 M) and 1 μL Glycogen (20 μg/μL) and precipitated with 600 μL of pre-chilled ethanol.

For sequencing, RNA sequencing libraries were constructed using the Stranded mRNA-seq Lib Prep Module for Illumina (RK20349, Abclonal) according to the manufacturer’s instructions. mRNA strand-specific library sequencing was performed at GenePlus (Beijing, China) using double-index libraries on the DNBSEQ-T7 platform in paired-end 150-bp mode.

#### Processing and alignment of RIP-seq datasets

Fastq files were checked for the reads quality with FastQC (v0.11.9). Raw reads were trimmed with trim galore (v0.6.4_dev, --paired) to remove low-quality bases and adapters from read pairs. BBDuk algorithm of BBMap was then used for removing rRNA-derived reads (k=27) matching the deposited ribosomal RNA sequencing (SILVA 128). For alignment, quality checked reads were aligned to the reference genome (hg38) using hisat2 (v2.1.0)^85^ with parameter “-no-mixed, -no-discordant, --rna-strandness RF”. Alignment files were then converted to bam format, sorted and indexed with samtools (v1.12). Reads for each reference gene (gencode 31) were quantified with featureCounts (v1.6.4, -p -Q 10 -s 2)^86^. Differential expression analysis was performed using DESeq2 (v1.28.1)^87^ to obtain enriched genes.

#### Validation of RNA-binding proteins by OOPS

OOPS was performed to validate the RNA binding ability of HMGCL and PHKG2. HEK293T cells were cultured in 10-cm-diameter dishes until reached a maximum of 90% of confluence. The cells were washed twice with PBS, and the supernatant was removed by pipetting. For the non-crosslinked controls, cells were immediately lysed by scraping in 1mL of acidic guanidinium-thiocyanate-phenol (TRIzol, Thermo Fisher Scientific), and the resulting homogenate was transferred to a new tube. For crosslinked samples, UV cross-linking was performed on PBS-washed cells by UV irradiation at 254 nm for 400mJ/cm² (Scientz 03-II Ultraviolet Crosslinker; Scientz). Immediately after crosslinking, the cells were scraped in 1mL TRIzol, and the homogenized lysate was transferred to a new tube and incubated at room temperature (RT) for 5 minutes to dissociate unstable RNA–protein interactions. Each of the two groups were equally divided into two fresh tubes, designated as RNase-group and RNase-free group.

For biphasic extraction, 100 µL of chloroform were supplemented, the phases were vortexed, and the sample was centrifuged for 15 minutes at 16,000 × g at 4°C. The upper aqueous phase and the lower organic phase were discarded, and the interface was further purified by performing an additional biphasic extraction with 1mL TRIzol and 200µL chloroform. The interface was then precipitated by addition nine volumes of methanol and pelleted by centrifugation at 16,000 × g for 10 minutes. The precipitated interface was resuspended by careful sonication in 100 µL of 100 mM triethylammonium bicarbonate (TEAB) and 1% SDS. The samples from RNase-treated group were incubated at 95 °C for 5 minutes, cooled on ice, and digested with 2 µL of RNase cocktail (Thermo Fisher Scientific) for 4h at 37°C. The released proteins in both the RNase-free and RNase-treated groups were finally extracted with 1mL TRIzol and 200 µL chloroform. The organic phase was transferred to a fresh tube and precipitated with four volumes of 100% ethanol. The pellet was solved in 0.4% SDS-PBS for further Western blot analysis.

#### Immunofluorescence staining and fluorescence microscopy

For TurboID labeling, HEK293T cells were seeded on 7 × 7-mm glass coverslips in 24-well plates and transiently transfected with 2 μg of TurboID-NES or TurboID-NLS for 24 hours. Cells were then treated with or without 50 μM biotin for 30 minutes. After the labeling, cells were washed twice with 1 mL of ice-cold PBS. The cells were then fixed with 4% paraformaldehyde for 15 minutes at room temperature and washed twice with 1 mL of PBS. Next, the cells were permeabilized using 0.5% Triton X-100 (diluted in PBS) for 15 minutes at room temperature and washed three times with 1 mL of PBS. The cells were blocked with 5% BSA dissolved in PBS for 1 hour. The coverslips were then incubated with primary antibodies against the V5-tag (1:200 dilution in PBS with 5% BSA, Thermo Fisher Scientific) overnight at 4°C. Afterward, the coverslips were washed three times in PBS, with each wash lasting 5 minutes, and then incubated with secondary antibodies conjugated to AlexaFluor-488 (1:200 dilution in PBS with 5% BSA, YEASEN) and streptavidin conjugated to AlexaFluor647 (1:200 dilution in PBS with 5% BSA, YEASEN) for 1 hour. The coverslips were washed five times in PBST (0.1% Tween-20 in PBS). Finally, the coverslips were incubated with DAPI (1:2000 dilution in PBST, Beyotime) for 10 minutes and washed three times.

To observe the location of SGK3, U2OS cells that stably express SGK3^WT^-EGFP cells or SGK3^DM^-EGFP were seeded in 20 mm glass confocal dish. For transfection of CASC15, 1 μg pcDNA 3.1 CASC15 was transfected with 1.2 μL lipo8000 in 6 well-plates, after 24 hours cells were digested by trypsin and seeded in 20 mm glass confocal dish. For IGF-1 stimulation, cells were treated with 100 ng/ml of IGF-1 for 20 minutes. Cells were washed twice with 1 mL of ice-cold PBS. Cells were fixed with 4% paraformaldehyde for 15 minutes at room temperature and washed twice with 1 mL of PBS. Cells were then permeabilized with 0.5% Triton X-100 (diluted in PBS) for 15 minutes at room temperature and washed three times with 1 mL of PBS three times. Cells were blocked in 5% BSA dissolved in PBS for 1 hour. The confocal dishes were incubated with primary antibody against EEA1 (1:100 dilution in PBS with 5% BSA) overnight at 4 °C. Cells were washed three times in PBS with 5 minutes for each wash then incubated in secondary antibodies conjugated to mouse AlexaFluor-647 (1:200 dilution in PBS with 5% BSA) for 1 hour. Cells were washed five times in PBST (0.1% Tween-20 in PBS). Dishes were incubated with DAPI (1:200 dilution in PBST) for 10 minutes then were washed for three times. Immunofluorescence images were taken using an AXR-NSPARC Super-Resolution Confocal Microscope.

To observe the nanoclusters of csRRPs on cell surface, HeLa cells were cultured on a confocal dish with glass bottom at the confluence of 25%, 24-36h in advance of treatment. To conduct cell surface immunostaining, anti-JAK1, anti-RO60, anti-YTHDC2, and rabbit IgG isotype were premixed in 0.5% BSA in PBS with goat anti-rabbit AF488 with molar ratio of 2:1 on ice for 1h. HeLa cells on confocal dishes were first washed twice with PBS, and then fixed with 4% PFA for 15min. After washing three times with PBS, cells were blocked with 1% BSA in PBS on ice for 30min. Then the mixture of antibody is added to the cells and incubated on ice for 1h. After another 3 washes with PBS, the cells were stained with Hoechst33342(1:5000) and WGA-Rhodamine(1:2000) at room temperature for 20min. Immunofluorescence images were taken using an AXR-NSPARC Super-Resolution Confocal Microscope.

#### RT-qPCR

For the validation of SGK3 CASC15 interaction by RIP-qPCR (Figure 3C), MCF7 cells were cultured in 6 cm dish and transiently transfected with 1 μg of pcDNA 3.1 CASC15 for 12 hours. After crosslinking under 400 mJ/cm^2^ by 254 nm UV light, cells were lysed. 40 μL of Protein G beads were washed 3 times with 450 μL of lysis buffer (50 mM Tris-HCl, pH 7.4, 100 mM NaCl, 1% NP-40, 0.1% SDS, 0.5% sodium deoxycholate, 1% EDTA-free protease inhibitor cocktail in DEPC H2O). Protein G beads were pre-conjugated with antibodies prior to immunoprecipitation with 200 μL of lysis buffer containing 2.5 μg of anti-SGK3 (Proteintech) with rotation at room temperature for 1 hour. After washing by lysis buffer for 3 times, the beads were then added into the cell lysates with rotation at 4 °C for 4 hours. Then following the same procedure described under “RIP-seq”. To assess the binding of wild-type SGK3 and the double-mutant to CASC15 by RIP-qPCR (Figure 3F), MCF7 stably expressing SGK3^WT^-EGFP and SGK3^DM^-EGFP cells were cultured in 6 cm dishes and transiently transfected with 1 μg of pcDNA 3.1 CASC15 for 12 hours. After crosslinking under 400 mJ/cm^2^ by 254 nm UV light, cells were lysed. 40 μL of anti-GFP beads were washed 3 times with 450 μL of lysis buffer (50 mM Tris-HCl, pH 7.4, 100 mM NaCl, 1% NP-40, 0.1% SDS, 0.5% sodium deoxycholate, 1% EDTA-free protease inhibitor cocktail in DEPC H2O). The beads were then resuspended in cell lysates with rotation at 4°C for 4 hours. Then following the same procedure described under “RIP-seq”. RNA samples were reverse transcribed using the HiScript III 1st Strand cDNA Synthesis Kit (R312, Vazyme) according to the manufacturer’s instructions. CASC15, SGK3 were analyzed by real-time quantitative PCR using the CFX96 instrument (Bio-Rad) and SYBR qPCR Master Mix (Q321, Vazyme). Normalized Ct values were calculated using the ΔΔCt method, and error bars represent the standard error of the mean (n = 3).

The primers used for RIP-qPCR assays:

CASC15-qPCR-F: CGCCGGGGTATCTCCCTCTCG
CASC15-qPCR-R: CATTTCCCCCGCTGCAGTCCA
SGK3-qPCR-F: ATGGCCCTGAAGATTCCTGC
SGK3-qPCR-R: ACGGTCCCAGGTTGATGTTC

#### RBR-ID

To identify the RNA binding sites of SGK3 (Figure 3D), RBR-ID was performed as previously described with modifications^65^. MCF7 cells were transiently transfected with 5 μg of Flag-SGK3 for 24 hours, while another set of MCF7 cells, which were not transfected with the plasmid, served as the negative control. The cells were washed twice with PBS, then re-coated with 5 mL of ice-cold PBS containing an EDTA-free protease inhibitor cocktail. For crosslinking, the plates (with lids removed) were placed on ice and crosslinked with 400 mJ/cm² UV light. After crosslinking, cells were harvested and lysed with Flag lysis buffer (50 mM Tris–HCl, pH 8.0, 137 mM NaCl, 1% Triton X-100, 1% sarkosyl, 1% glycerol, 1 mM Na3VO4, 1 mM DTT, 1 mM PMSF, and 1% protease cocktail), followed by sonication on ice. The soluble lysates were collected after ultracentrifugation at 20,000 × g for 20 minutes at 4°C. The protein concentration of the supernatant was determined using the BCA protein assay kit (Pierce). All protein concentrations were normalized to the same level. The lysates were incubated overnight with 20 μL of anti-Flag M2 affinity gel beads (Sigma) at 4°C with gentle rotation. The resulting beads were washed twice with BC200 buffer (200 mM NaCl, 20 mM Tris, pH 7.3, 20% glycerol, 0.2% NP-40) and three times with BC100 buffer (100 mM NaCl, 20 mM Tris, pH 7.3, 20% glycerol, 0.1% NP-40). The enriched proteins were eluted by boiling the beads in 40 μL of 1× protein loading buffer at 95°C for 10 minutes. The supernatant was separated by 10% SDS-PAGE and subjected to in-gel trypsin digestion for subsequent LC-MS/MS analysis.

#### Immunoprecipitation

For detecting SGK3 phosphorylation in Figure3J, HEK293T SGK3^WT^-EGFP cells were cultured in 6-well plates and transfected with 1 μg pcDNA3.1 CASC15 for 24 hours, then the IGF-1 treated groups were treated with 100 ng/mL IGF-1 for 20 min, cells were collected, washed twice with PBS, and then centrifuged at 3,000 rpm for 4 minutes. The cells were lysed in 100 μL BC100 buffer containing EDTA-free protease inhibitor cocktail for 1 hour with rotation at 4 °C. The lysates were then centrifuged at 20,000 × g for 20 minutes. The protein concentration in the supernatant was measured using a BCA protein assay kit (Pierce). Anti-GFP Affinity beads were washed five times with BC100 buffer. The lysates were incubated with anti-GFP Affinity beads overnight at 4 °C with rotation. The supernatants were discarded, and the beads were washed five times with BC100 buffer. Proteins were eluted from the beads by incubating with 50 μL of elution buffer (0.1 M glycine HCl, pH 3.0) for 5 minutes and neutralizing with 5 μL 1 M Tris-HCl, pH8.0 neutralization buffer. The samples were then processed according to the Western blot protocol.

#### Protein purification

Purification of Flag-SGK3 PX domain (1-126) was shown in FigureS3D, pGEX-6P-1 Flag-SGK3^PX^ plasmid was constructed. Plasmids were then transformed into competent E. coli BL21 (DE3) expression strain via the heat shock method, followed by antibiotic selection on LB agar plates containing ampicillin. Single clone was selected in 5 mL LB medium containing ampicillin in 37 ℃ incubating shaker overnight. The overnight-cultured bacterial suspension was inoculated into 1 L of LB medium in incubating shaker at 37 ℃ for 3 hours. 0.5 mM IPTG was added, and induced expression was carried out at 16 °C for 16 hours before collected.

The cell pellet was resuspended in lysis buffer (50 mM Tris pH 8.0, 300 mM NaCl, 0.5 mM TCEP) and was disrupted using a high-pressure cell disruptor. Then the lysate was centrifuged at 15000 rpm for 40 min, the supernatant was collected and incubated with Glutathione Sepharose 4B resin at 4 ℃ for 2 hours. Then the Glutathione Sepharose 4B resin was washed by protein buffer and was incubated with GST-PreScission Protease at 4 ℃ overnight. The flowthrough was collected and concentrated to 2 mL and further purified by a Superdex 200 10/300 GL size-exclusion column (GE Healthcare) pre-equilibrated with PBS buffer, attached to an AKTA pure (GE Healthcare). The purified protein was concentrated to 1 mg/ml.

#### Differential Scanning Fluorimetry

The Differential Scanning Fluorimetry assay was performed with a BioRad 96 system (Bio-Rad, CFx 96 Touch). A mixture containing PBS, 2 μM recombinant Flag-SGK3^PX^ protein, SYPRO Orange dye, Ptdlns(3)P and CASC15 RNA (different concentrations were used in different preparations) was heated from 25 °C to 95 °C at a 1% increment. The fluorescence intensity was detected at 492 nm excitation and 610 nm emission wavelengths. The normalized *Tm* data were calculated by OriginPro 2021.

#### Lipid overlay assay

For exploring the effect of CASC15 RNA in modulating Ptdlns(3)P and SGK3 PX domain binding in Figure 3H, lipids Ptdlns(3)P and Ptdlns(4)P (different concentrations were used) were doted on nitrocellulose membrane and dry out for 30 minutes. The 3 μM recombinant Flag-SGK3^PX^ protein was incubated with or without 3 μM CASC15 for 1 hour and crosslinked by 254 nm UV light. Then incubated proteins were diluted in blocking buffer (50 mM Tris-HCl pH7.5, NaCl 150 mM, 0.1 % Tween 20, 0.4% BSA) to make 10 nM concentration solutions. The protein solutions were added to the lipid doted nitrocellulose membrane, incubated at 4 ℃ overnight. Then the nitrocellulose membrane was washed 3 times by TBST, incubated with anti-Flag HRP (1:1000) at room temperature for 1 hour, washed 3 times by TBST. Then developed Clarity Western ECL Blotting Substrates (Bio-Rad) and imaged on the Tanon 5200 Multi System (Tanon).

#### Flow cytometry

For flow cytometry in Figure 7H, cells were cultured as above and gently lifted with accutase (gibco) for 3 minutes at 37°C, quenched with DMEM, and then counted. For each condition, 50,000 cells were used and blocked with Human TruStain FcX (Fc block, BioLegend) in FACS buffer (0.5% BSA in 1× PBS) for 30 minutes on ice, and then were incubated with premixed antibody solution on ice for another 30 minutes. For the premixed antibody, anti-RO60, anti-YTHDC2, anti-JAK1, and rabbit isotype were separately mixed with goat anti-rabbit AF488 antibody with the molar ratio of 2:1 in FACS buffer on ice for 30min. After incubation, the cells were washed twice with FACS buffer and flow cytometry with FITC channel was conducted with BD LSRFortessa SORP. Data collected was processed and analyzed with FlowJo.

#### cell surface RNA FISH

For cell surface RNA FISH experiments in Figure S7H and Figure S7I, HeLa cells were cultured on a confocal dish with glass bottom at the confluence of 25%, 24-36h in advance of treatment. HeLa cells on confocal dishes were first washed twice with PBS, and then fixed with 4% PFA for 15min. Cells were afterwards washed sequentially with three times of PBS and three times of 10% formamide in 2×SSC buffer. Then the cells were incubated with 1μM 1^st^ probe in 10% formamide, 2×SSC buffer, and 10% dextran sulfate at 37℃ overnight. The next day, cells were. washed three times with pre-warmed 30% formamide in 2×SSC buffer, 10 min per wash, at 37°C with gentle rocking, then incubated with 1μM 2^nd^ probe in 30% formamide and 2×SSC buffer at 37°C for 1h. Then the cells were washed 4 times with PBS, 15min per wash. After that, the cells can also be co-stained with csRRP antibodies as described in “Immunofluorescence staining and fluorescence microscopy”, or directly stained with Hoechst33342 to label the nucleus. For the group with RNase treatment, 0.02mg/mL RNase A in Hanks buffer were used to cleave the RNAs on cell surface at 37°C for 20min before fixation with PFA.

Y5 RNA 1^st^ probe: GGGAGACAATGTTAAATCTTTACATCATCATACATCATCATACATCATCAT

Y5 RNA 2^nd^ probe: ATGATGATGTTTTTTTTT-TAMRA

### QUANTIFICATION AND STATISTICAL ANALYSIS

For comparison between two groups, p-values were determined using two-tailed Student’s t tests, \**p* < 0.05; \*\**p* < 0.01; \*\*\**p* < 0.001; \*\*\*\**p* < 0.0001. N.S. not significant. For all box plots, p-values were calculated with Wilcoxon rank-sum test, \**p* < 0.05; \*\**p* < 0.01; \*\*\**p* < 0.001; \*\*\*\**p* < 0.0001. Error bars represent means ± SD. R² was computed by performing linear regression analysis, where the square of the correlation coefficient (Pearson’s r) with 95 % confidence interval was determined

